# Heterosis is a systemic property emerging from nonlinear genotype-phenotype relationships: evidence from *in vitro* genetics and computer simulations

**DOI:** 10.1101/170902

**Authors:** Julie B. Fiévet, Thibault Nidelet, Christine Dillmann, Dominique de Vienne

## Abstract

Heterosis, the superiority of hybrids over their parents for quantitative traits, represents a crucial issue in plant and animal breeding. Heterosis has given rise to countless genetic, genomic and molecular studies, but has rarely been investigated from the point of view of systems biology. We hypothesized that heterosis is an emergent property of living systems resulting from frequent concave relationships between genotypic variables and phenotypes, or between different phenotypic levels. We chose the enzyme-flux relationship as a model of the concave genotype-phenotype (GP) relationship, and showed that heterosis can be easily created in the laboratory. First, we reconstituted *in vitro* the upper part of glycolysis. We simulated genetic variability of enzyme activity by varying enzyme concentrations in test tubes. Mixing the content of “parental” tubes resulted in “hybrids”, whose fluxes were compared to the parental fluxes. Frequent heterotic fluxes were observed, under conditions that were determined analytically and confirmed by computer simulation. Second, to test this model in a more realistic situation, we modeled the glycolysis/fermentation network in yeast by considering one input flux, glucose, and two output fluxes, glycerol and acetaldehyde. We simulated genetic variability by randomly drawing parental enzyme concentrations under various conditions, and computed the parental and hybrid fluxes using a system of differential equations. Again we found that a majority of hybrids exhibited positive heterosis for metabolic fluxes. Cases of negative heterosis were due to local convexity between certain enzyme concentrations and fluxes. In both approaches, heterosis was maximized when the parents were phenotypically close *and* when the distributions of parental enzyme concentrations were contrasted and constrained. These conclusions are not restricted to metabolic systems: they only depend on the concavity of the GP relationship, which is commonly observed at various levels of the phenotypic hierarchy, and could account for the pervasiveness of heterosis.

## Introduction

The relationship between genetic polymorphism and phenotypic variation is a major issue in many branches of biology. It is usually referred to as the genotype-phenotype (GP) relationship, GP map or GP correlation. Strictly speaking, the genotype corresponds to the DNA level, whereas the phenotype can correspond to any trait observed or measured at any organizational level from transcription rate to fitness. In fact, following Wright [1], the GP relationship can also refer to the relationship between a genetically variable phenotypic level (e.g. the activity of an enzyme) and a more integrated phenotypic level (e.g. a flux through a pathway).

Almost inevitably, the relationship between parameters at different organizational levels is nonlinear, because the cellular processes that shape biological functions and structures are highly intertwined. The kinetics of biochemical and molecular reactions, transport and signaling mechanisms, positive and negative feedbacks, growth kinetics, etc., are intrinsically nonlinear. Thus, from allosteric regulation to fitness landscapes, from genetic regulatory circuits to developmental processes, there are myriads of examples showing complex relationships between genotype and phenotypes, or between phenotypic levels [2, 3]. These GP relationships display a large diversity of shapes [4–6], but two prevail to a large extent:

i. The sigmoidal relationship (Fig 1A). Cooperativity and synergy underlie such behavior [7, 8], which is classically modeled using Hill functions. For instance in Drosophila, transcription of the *hunchback* gene in response to the gradient of Bicoid transcription factor concentration follows a typical S-shaped curve [9]. Examples at other phenotypic levels exist, such as the relationship between the expression level of various genes and growth rate, a proxy for fitness, in *Escherichia coli* [10, 11] and in *Saccharomyces cerevisiae* [12].
ii. The concave relationship (Fig 1B), the archetype of which is the enzyme-flux relationship. The long-standing metabolic control theory (MCT; reviewed in [13]) is probably the most elaborated corpus to describe this particular, but essential, case of genotype-phenotype relationship. Both theoretical developments and experimental observations show that increasing from zero the value of an enzyme parameter usually results first in an increase of the flux value, then in attenuation of the response and finally in saturation. The phenotypic response resembles a rectangular hyperbola, and at the asymptotic upper limit there is robustness of the phenotype towards genotypic variation.

**Fig. 1.**
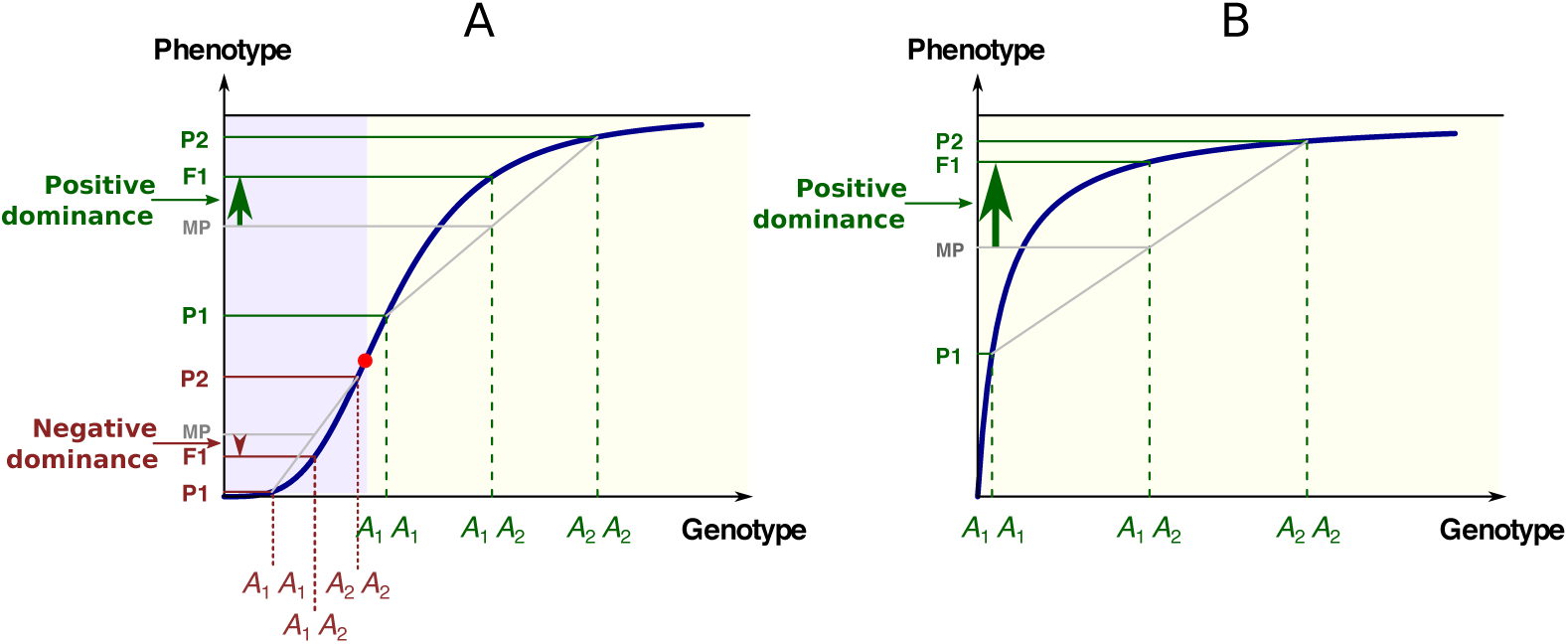
The two most common types of GP relationships with possible inheritance. A: The S-shaped relationship. There is convexity below the abscissa of the inflection point (red point) and concavity above. The three phenotypes P1, F1 and P2 are associated with the three genotypes *A*_1_*A*_1_, *A*_1_*A*_2_ and *A*_2_*A*_2_, respectively. MP: mid-parental value. In the convex region of the curve (mauve background), the low allele is dominant over the high allele (negative dominance), while in the concave region of the curve (yellow background) the high allele is dominant over the low allele (positive dominance). B: The hyperbolic concave relationship. Whatever the genotypic values, there is positive dominance. In all cases there is additive inheritance of the genotypic parameters.

Such concave relationships are common at other organizational levels. For instance, it is observed during transcriptional regulation when transcription factors do not bind to DNA cooperatively [14]. In yeast, the rate of protein synthesis in response to variation in abundance of some twenty different translation factors is concave in most cases [15], and can be modeled using a deterministic model [16, 17]. Other examples show the effects of variable gene/protein expression on the fitness of an organism. Izard et al. [18], by engineering the control of RNA polymerase expression in *E. coli*, consistently observed a saturation curve for growth rate in diverse media. In mitochondrial diseases, the relationship between the proportion of mtDNA carrying pathogenic mutations and various cellular responses – mitochondrial protein synthesis rate, respiratory chain complex activity, mitochondrial respiration or ATP synthesis and growth rate – is concave, with a large plateau. Thus an effect will only be detectable above a threshold value of the proportion of deleterious mtDNA (reviewed in [19]). In Parkinson’s disease, the symptoms do not begin to appear until about 80% of neurons in the substantia nigra, which produce dopamine, have died. This is consistent with the hyperbola-shaped relationship between the proportion of alive neurons and the concentration of extra-cellular dopamine [20]. In a wide range of species, allometry also provides relevant examples of non-linear relationships, such as between organismal mass and whole-organism metabolic rate, but also life span, tree trunk diameter, blood circulation time, etc. [21]. Allometric equations are power laws whose scaling exponents are lower than 1, resulting in concave relationships.

In the above-mentioned studies, only one genotypic (or phenotypic) parameter is assumed to be variable. In fact, most integrated phenotypic traits are polygenic, i.e. they are potentially affected by a very large number of parameters, giving rise to complex and rugged phenotypic landscapes. However, from the previous considerations, one can expect global concavity of these multi-dimensional surface responses. Indeed, within the MCT framework, Dykhuizen et al. [22] showed that the fitness response of *E. coli* to variation of both *β*-galactoside permease and *β*-galactosidase activity is a two-dimensional hyperbolic-like surface. Similarly, Nijhout et al. [23] analyzed quantitatively the pairwise effects of various components of the MAPK signaling cascade on the MAPK-PP output and observed largely concave response surfaces with saturation. McLean [24] also found a concave relationship between growth rate and the joint variation of transcription and translation mediated by streptomycin and rifampicin, respectively, in *Pseudomonas aeruginosa*. Another interesting example is the allometric relationship described in a population of recombinant inbred lines of *Arabidopsis thaliana* between, on the one hand, photosynthetic rate and, on the other hand, dry mass per area and age at flowering [25].

Whether the phenotypic response is sigmoidal or concave, there is often a horizontal asymptote which accounts for the strong asymmetry of the phenotypic response, depending on whether the value of the genotypic variable is increased or decreased. Thus, in metabolic engineering it is usually much easier to decrease than to increase flux and metabolite concentrations [26]. At the genome-wide level, this asymmetry was recognized early on through the classical work of Lindsley et al. [27] on the effects of segmental aneuploidy in Drosophila. Using *Y* -autosome translocations with defined breakpoints, the authors produced a large number of small duplications and deficiencies spanning chromosomes 2 and 3. Examining the phenotypic effects of having the same chromosomal region present in one, two or three copies, the authors found that monosomy had a much greater effect on viability than trisomy, revealing a strong asymmetry of the response to gene dosage. “The pervasive robustness in biological systems” [6], which has been observed at all phenotypic levels (transcript, protein and metabolic abundance, metabolic flux, chemotaxis, cell cycle period, cell fate patterning, cell survival, etc.), most likely stems from this saturation effect. In some cases the mechanisms of robustness have been unraveled [28–30], and a widespread link is thought to exist between robustness and network complexity [31, 32], even though robustness could be partly adaptive [33].

In addition to robustness, the “diminishing return” curves could account for various fundamental and apparently unrelated observations. The best-known example is the dominance of “high” over “low” alleles for metabolic genes, which was first pointed out by Wright [1], then formalized in detail by Kacser & Burns [34] in a seminal paper. Due to concavity, the phenotypic value of the heterozygote exceeds the homozygote mean value, and alleles with large deleterious homozygous effects are more recessive than alleles of weaker effect (Fig 1). The metabolic concave relationship also has major implications regarding the evolution of selective neutrality [35], the variability of enzyme activity in populations under mutation-selection balance [36], the relationship between flux and fitness [22], the epistasis between deleterious mutations [37], the response to directional selection [38], the distribution of QTL effects [39, 40], the dynamics of retention/loss of metabolic genes after whole genome duplications [41] and the evolution of genes in branched pathways [42].

The model of positive dominance of Wright, Kacser and Burns is not restricted to metabolic genes, since it is valid whenever there is concavity of the GP relationship. In the case of a sigmoidal response, dominance of deleterious mutations may occur, depending on the allelic values relative to the inflection point of the curve [43, 44] (Fig 1A). It would be quite a laborious task to try to estimate the respective proportions of positive and negative dominance in the literature. Nevertheless, the fact remains that there is a marked bias towards the dominance of “high” alleles. As early as 1928, Fisher [45] noted that, in Drosophila, 208 out of 221 autosomal or sex-linked deleterious mutations were recessive. Similar proportions have been observed in all species studied to date, including in artificial diploids of the normally haploid *Chlamydomonas* [46]. More recently, genome-wide estimates from collections of deletion strains in yeast have been published [47, 48]. Whereas 891 genes (20% of the genome) contributed to slow growth as homozygotes, only 184 (3% of the genome) were haploinsufficient [48]. The mean values of the dominance coefficients were found to be *h* = 0.25 or less, which means that null alleles were on average recessive [49, 50]. Finally, predominant recessivity of deleterious mutations is fully consistent with common observations in population genetics, such as inbreeding depression [51].

In this paper, we show that generalizing this dominance model to the multilocus case directly leads to a simple model of heterosis, i.e. a model that accounts for the common superiority of hybrids over their parents for quantitative traits [52, 53]. Described more than 250 years ago (cited in [54]), hybrid vigor is of great importance for food production because it affects complex traits such as growth rate, biomass, fertility, disease resistance, drought tolerance, etc. In crops, it is currently used to derive hybrid varieties with exceptional performances. For instance in maize, the hybrid grain yield can exceed by more than 100 % the mean parental yield [55, 56]. In rice, heterosis is about 20 % for grain yield and about 10 % for plant height [57]. In livestock and farms animals, such as cattle, swine, sheep and poultry, heterosis is also present and commonly exploited [58, 59]. This phenomenon is not restricted to species of agronomic interest. In the model plant *A. thaliana*, there is heterosis for biomass, rosette diameter and flowering date [60–62]. In animals, it has been described for diverse traits in drosophila [63], frog [64], Pacific oyster [65], owned dogs [66], etc. Heterosis has also been found in microorganisms, for instance for growth rate in *Neurospora* [67] and *S. cerevisiae* [68]. In the latter case it may be spectacular, with growth rate one order of magnitude higher in the hybrid than in the best parent.

Theoretical models of heterosis have been proposed based on the functioning of gene circuits [69] or metabolic systems [70,71]. In the latter case the authors unraveled the links between heterosis, dominance and epistasis within the MCT framework. In support of this model, we show here that heterosis can easily be created in the laboratory. We relied on two complementary approaches: (i) Test-tube genetic experiments. We crossed *in vitro* “parents” that differed in the concentration of four successive glycolysis enzymes, and obtained “hybrids” displaying heterosis for flux through the pathway; (ii) *In silico* genetics. We crossed parents that differed in the concentration of 11 enzymes from the carbon metabolism network in *S. cerevisiae*, and we computed the input flux and two output fluxes in parents and hybrids using a system of differential equations. Heterotic fluxes were frequently observed. The conditions for having heterosis were analyzed using computer simulations and mathematically using the theory of concave functions. The phenotypic proximity of parents, together with contrasted and constrained parental enzyme concentrations, proved to be reliable predictors of heterosis. These conclusions are qualitatively valid beyond metabolic networks because they only depend on the shape of the GP relationship. Thus the so-called “mysterious” phenomenon of heterosis appears to be a systemic property emerging from the nonlinearity that exists at various levels of the phenotypic hierarchy.

## Materials and methods

### Theoretical background

#### Heterosis indices

Let the relationship between a phenotypic variable *z* and a series of *n* genotypic variables *x*_*j*_:

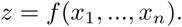

Consider two individuals P1 and P2 with distributions of genotypic values {*x*_*i*1_}_*i*=1,…,*n*_ and {*x*_*i*2_}_*i*=1,…,*n*_, respectively, and their corresponding phenotypic values:

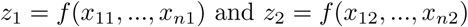

The hybrid P1‪P2 has the genotypic distribution {*x*_*i*,1*∗*2_}_*i*=1,…,*n*_ and the phenotypic value:

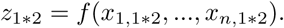

When comparing *z*_1*∗*2_ with the mean of *z*_1_ and *z*_2_, heterosis can be observed. In its most common definition, heterosis corresponds to a positive departure of the hybrid value from the mean parental value for quantitative traits. However negative heterosis also exists, and thus five situations of inheritance can be defined [72–74] (S1 Fig). Following our notations, we have:

Best-Parent Heterosis (BPH): *z*_1*∗*2_ *>* max(*z*_1_*,z*_2_)
positive Mid-Parent Heterosis (+MPH): *z*_1*∗*2_ *∈*](*z*_1_ + *z*_2_)*/*2, max(*z*_1_,*z*_2_)]
Additivity (ADD): *z*_1*∗*2_ = (*z*_1_ + *z*_2_)*/*2
negative Mid-Parent Heterosis (–MPH): *z*_1*∗*2_ *∈* [min(*z*_1_*,z*_2_), (*z*_1_ + *z*_2_)*/*2)[
Worst-Parent Heterosis (WPH): *z*_1*∗*2_ *<* min(*z*_1_*,z*_2_).

A common index of heterosis, the potence ratio [75]:

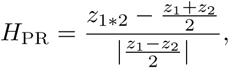

directly gives the type of inheritance:

If *H*_PR_ *>* 1, there is BPH.
If *H*_PR_ *∈*]0, 1], there is +MPH.
If *H*_PR_ = 0, there is ADD.
If *H*_PR_ *∈* [−1, 0[, there is –MPH.
If *H*_PR_ *< −*1, there is WPH.

We frequently used in this study the index of best-parent heterosis:

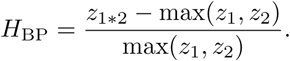

If *H*_BP_ *>* 0, there is BPH.
If *H*_BP_ *≤* 0, there can be +MPH, ADD, –MPH or WPH.

The mean of the positive *H*_BP_ values was noted 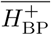.

We also defined an index of worst-parent heterosis:

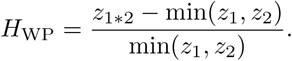

If *H*_WP_ *<* 0, there is WPH.
If *H*_WP_ *≥* 0, there can be –MPH, ADD, +MPH or BPH.

The mean of the negative *H*_WP_ values was noted 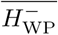.

Finally, we used in some instances the index of mid-parent heterosis:

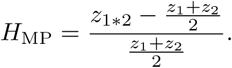

If *H*_MP_ *>* 1, there is BPH or +MPH.
If *H*_MP_ = 0, there is ADD.
If *H*_MP_ *<* 1, there is –MPH or WPH.

Note that in this context, various authors wrongly refer to heterosis as overdominance, complete dominance, partial dominance, and use various circumlocutions for negative heterosis. Strictly speaking, the words “dominance” and overdominance only apply to monogenic traits.

#### The concave (resp. convex) genotype-phenotype relationship results in positive (resp. negative) heterosis

Consider 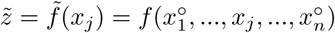, defined as the function obtained by fixing 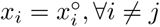, and letting *x*_*j*_ be variable. Assume that 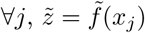 is strictly concave. Following the standard concavity argument, we can write:

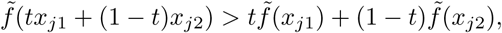

where *t ∈*]0, 1[. The additivity of genotypic variables corresponds to *t* = 1/2 for all *j*, so we have:

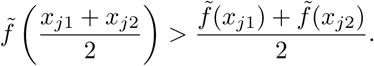

Hence

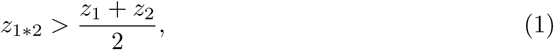

which means that +MPH or BPH necessarily holds, and that:

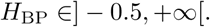

Note that if the parents have the same phenotypic value (*z*_1_ = *z*_2_), it comes from Eq 1 that there is necessarily BPH.

If 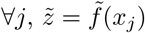 were strictly convex, Eq 1 would become:

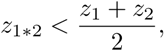

and there would be necessarily –MPH or WPH.

#### Application of these theoretical models to the case of the enzyme-flux relationship

Since Wright [1], the enzyme-flux relationship is considered the archetype of the concave genotype-phenotype (GP) relationship. Using a simple system of differential equations, he relied on the concavity of the GP curve to propose his famous model of dominance. Kacser & Burns [34] refined his approach using a formalism in which the relationship between enzyme concentrations and the flux through a metabolic pathway is described by a heuristic equation which is at the heart of the metabolic control theory [3, 13, 34]. Similarly, but more generally, Fiévet et al. [71, 76] expressed the relationship between the overall flux *J* (M^−1^.s^−1^) through a metabolic network and *n* enzyme concentrations {*E*_*j*_}_*j*=1,…,n_ in the following way:

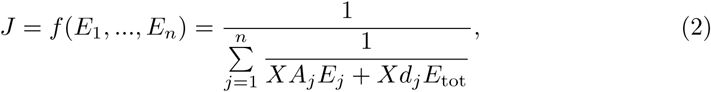

where *E*_*j*_ is the concentration of enzyme *j*, *X* is a positive constant, *A*_*j*_ and *d*_*j*_ are systemic parameters (unit M^−1^.s^−1^) that account for the kinetic behavior of enzyme *E*_*j*_ in the network, and *E*_tot_ is the total enzyme amount: 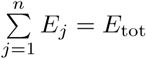. Because in the standard case *A*_*j*_ and *d*_*j*_ are positive, the flux is an ascending function of *E*_*j*_, for all *j*. We will consider that *A*_*j*_ and *d*_*j*_ are not genetically variable, so that variation in enzyme activity will only be due to the variability of the enzyme concentration *E*_*j*_. Nevertheless formal developments with variable *A*_*j*_’s would also be possible since *A*_*j*_ and *E*_*j*_ play a similar role in the relationship.

##### Flux heterosis

The variables and parameters of Eq 2 are all positive, and the function *J* = *f*(*E*_1_, …, *E*_*n*_) has a negative partial second derivative with respect to *E*_*j*_, for all *j*. Therefore the function is strictly concave, and the previous developments apply. In particular, consider two individuals P1 and P2 that differ in their distribution of enzyme concentrations, respectively {*E*_*j*1_}_*j*=1,…,*n*_ and {*E*_*j*2_}_*j*=1,…,*n*_. Assuming additive inheritance of enzyme concentrations, the distribution of enzyme concentrations of the P1*P2 hybrid is:

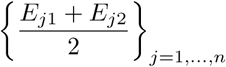

and its flux is:

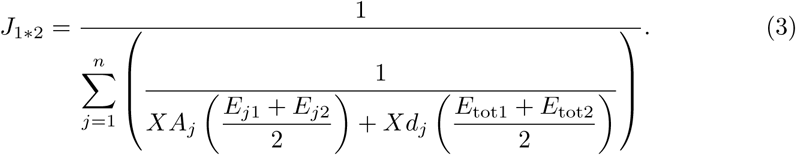

Due to concavity, we have for all *j*:

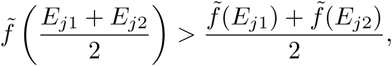

so

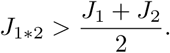

Thus either +MPH or BPH are expected in this GP relationship, and if the parents have the same flux value (*J*_1_ = *J*_2_) there is necessarily BPH.

##### Conditions for Best-Parent Heterosis

In the particular case where there is only one variable enzyme, BPH cannot occur, as can be shown from Eq 2. Otherwise, with *J*_2_ *> J*_1_, the condition for having BPH is *J*_1*∗*2_ *> J*_2_, i.e.:

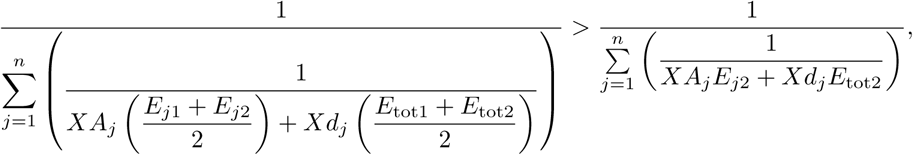

or

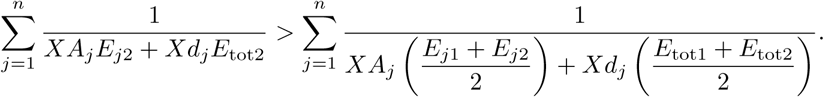

After rearrangements, we get:

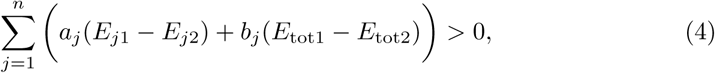

where

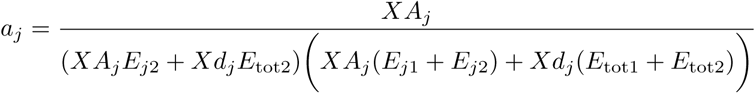

and

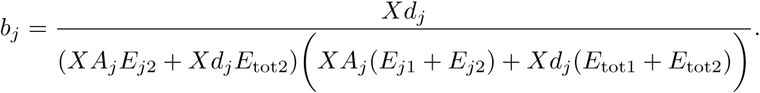

Note that *∀j, a*_*j*_ and *b*_*j*_ are positive.

If the total enzyme concentration does not vary between individuals due to global cellular constraints of space and energy, we have *E*_tot1_ = *E*_tot2_, and the condition for BPH is simplified:

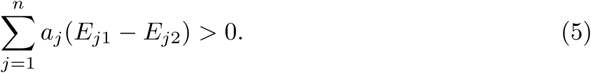

In this case, enzyme concentrations do not vary independently, and this covariation will increase heterosis because the concavity of the flux-enzyme relationship is higher. This can be shown by comparing the flux control coefficient 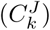 [34] to the combined response coefficient 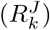 [77]. Both coefficients measure the sensitivity of the flux to variations in enzyme concentration *k*, but the former applies when concentrations vary independently whereas the latter applies when there is a constraint on the total enzyme concentration. Following [77], we have:

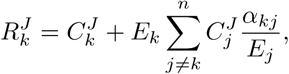

where 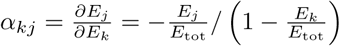 is the redistribution coefficient, i.e. the ratio of an infinitesimal change in concentration *E*_*j*_ to an infinitesimal change in concentration *E*_*k*_. As 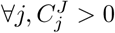 and *∀*(*k, j*), *α*_*kj*_ < 0, we have 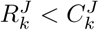, which means that constraint on *E*_tot_ increases concavity.

##### Optimal distribution of enzyme concentrations

The optimal distribution of enzyme concentrations is the one that maximizes the flux through the network for a given *E*_tot_ value. The relevant unit of measure here is g.L^−1^ and not M.L^−1^, because it is the total weight that matters in terms of energy and crowding in the cell. So we re-wrote Eq 2:

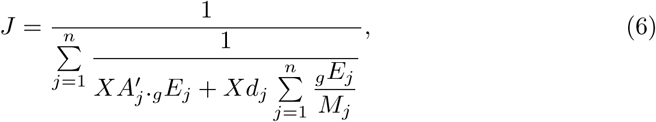

where 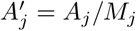, with *M*_*j*_ the molecular weight of enzyme *j*, and where _*g*_*E*_*j*_ is the concentration of enzyme *j* in g.L^−1^. The constraint is on 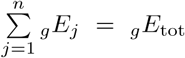, which is different from a constraint on 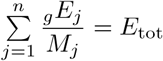. We derived the optimal distribution of enzyme concentrations, 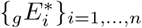, using the Lagrange multiplier method (see [76] for details):

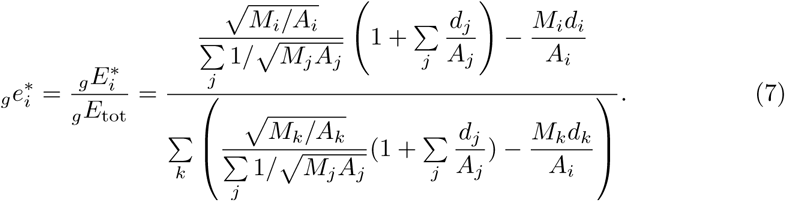

#### *In vitro* and *in silico* genetics

##### *In vitro* crosses

The *in vitro* system based on the upstream part of glycolysis is described in [76]. It goes from hexokinase (HK, E.C. 2.7.1.1) to glycerol 3-phosphate dehydrogenase (G3*P*DH, E.C. 1.1.1.8) (Fig 2). All assays were carried out at 25^*◦*^C in 50 mM Pipes buffer pH 7.5 containing 100 mM glucose, 100 mM KCl, 20 mM phosphocreatine, 3 mM NADH and 5 mM Mg-acetate. The reaction was started by the addition of 1 mM ATP. The ATP pool was kept constant via the regeneration reaction catalyzed by creatine phosphokinase. The high concentration of G3*P*DH (1 mM) allowed us to measure the flux through the pathway as the rate of NADH consumption at 390 nm, using an Uvikon 850 spectrophotometer.

**Fig. 2.**
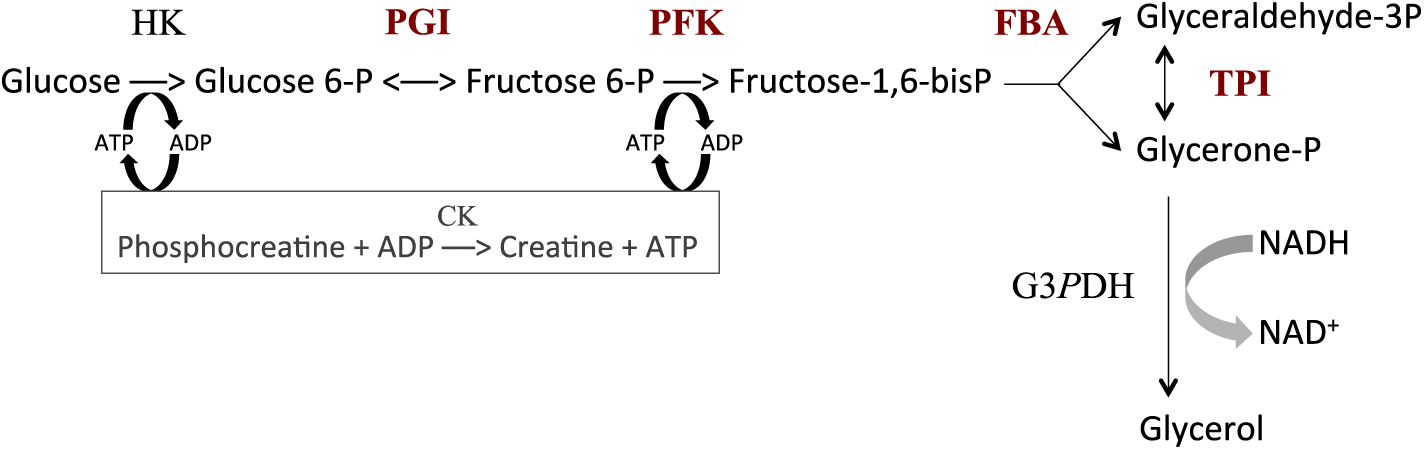
The upstream part of glycolysis reconstructed *in vitro*. HK, hexokinase (E.C. 2.7.1.1); PGI, phosphoglucose isomerase (E.C. 5.3.1.9); PFK, phosphofructokinase (E.C. 2.7.1.11); FBA, fructose-1,6-bisphosphate aldolase (E.C. 4.1.2.13); TPI, triosephosphate isomerase (E.C. 5.3.1.1); G3*P*DH, glycerol 3-phosphate dehydrogenase (E.C. 1.1.1.8); CK, creatine phosphokinase (E.C. 2.7.3.2). Variable enzymes are in red. Reaction rate was measured from the decrease in NADH concentration.

In order to mimic genetic variability in enzyme activity, we varied the concentrations of PGI, PFK, FBA and TPI in test tubes to create “parents”, each parent being defined by a particular vector of concentrations. HK concentration was fixed at 5.37 mg.L^−1^. The total enzyme concentration of the four variable enzymes in the parental tubes was fixed at 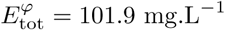, a value chosen from the physiological concentrations estimated in yeast strain S288C [78] (PGI: 9.1 mg.L^−1^, PFK: 10.4 mg.L^−1^, FBA: 60.1 mg.L^−1^ and TPI: 22.3 mg.L^−1^). To have parental genotypes covering a large range of enzyme concentrations, we varied the concentration of FBA (the most abundant enzyme) from 0 to 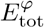, taking 10 values evenly distributed across this range (excluding of course the two extreme values). The proportions of the remaining enzymes (PGI, PFK and TPI) were drawn from beta distributions with shape parameter *α* = 1 and scale parameter 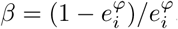, with 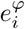 the physiological proportion of enzyme *i*. We thus created 58 parental distributions.

”Hybrids” were produced by mixing 1:1 the content of the parental tubes, which corresponds to additivity of enzyme concentrations. In order to have hybrids derived from a large range of parental values, we defined four classes of parental flux values (*<* 5 *µ*M.s^−1^, 5–8 *µ*M.s^−1^, 8–11 *µ*M.s^−1^ and *>* 11 *µ*M.s^−1^), and we performed “crosses” within and between classes. We created 61 hybrids using 36 different parents (S1 Table). For each of the 97 genotypes (61 hybrids + 36 parents), three flux measurements were performed, with the exception of two parents and four hybrids for which there were only two replicates (S2 Table).

Two statistical tests were performed: (i) a Student’s *t*-test to compare hybrid and mean parental fluxes; (ii) a Tukey’s test to classify observed inheritance as either BPH, +MPH, ADD, –MPH or WPH. For both, *p*-values were adjusted for multiple tests using the conservative Holm’s method.

##### Computer simulations of the *in vitro* system

We simulated the *in vitro* system using the parameter values published by Fiévet et al. [76], who estimated *XA_j_* and *Xd_j_* by hyperbolic fitting of the titration curves obtained by varying the concentration of each enzyme in turn. The values were *XA*_PGI_ = 499.4 s^−1^*, XA*_PFK_ = 115.5 s^−1^*, XA*_FBA_ = 22.5 s^−1^*, XA*_TPI_ = 22940 s^−1^*, Xd*_PGI_ = 0*, Xd*_PFK_ = 0*, Xd*_FBA_ = 0 and *Xd*_TPI_ = 21.9 s^−1^. Thus for any set of *E*_*j*_ values, i.e. for any virtual genotype, the flux value could be computed from the equation:

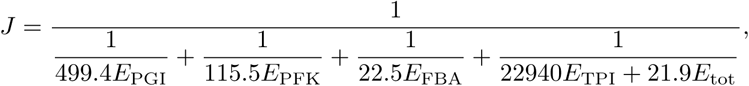

where *E*_tot_ = *E*_PGI_ + *E*_PFK_ + *E*_FBA_ + *E*_TPI_. Because enzyme concentrations in the simulations were in mg.L^−1^ and flux is expressed in *µ*M.s^−1^, we used the equation:

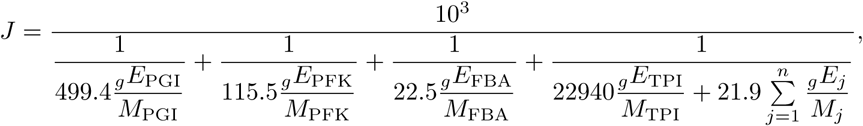

with the molecular masses *M*_PGI_ = 61128, *M*_PFK_ = 36000, *M*_FBA_ = 39211 and *M*_TPI_ = 26700.

We analyzed the effects of three factors on heterosis: (i) mean parental enzyme concentrations: either equidistributed or centered on their optima; (ii) constraint on total enzyme concentrations: either free *E*_tot_ or fixed *E*_tot_; (iii) changes in enzyme concentrations, by varying their coefficients of variation. For equidistributed enzyme concentrations, the means of the gamma distributions were *µ* = 25.475 mg.L^−1^ (i.e. 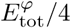) whatever the enzyme. For the optimum-centered distributions, the means were *µ* = 15.768 mg.L^−1^, *µ* = 25.162 mg.L^−1^, *µ* = 59.497 mg.L^−1^ and *µ* = 1.473 mg.L^−1^ for PGI, PFK, FBA and TPI, respectively, computed from Eq 7. In all cases we varied the coefficients of variation from *c*_v_ = 0.1 to *c*_v_ = 1.2, within the range of observed *c*_v_’s for these enzymes [79]. The shape and scale parameters of the gamma distributions were respectively 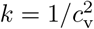 and 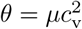.

To apply a strict constraint on the total amount of enzymes allocated to the system, we assumed that for each parent *i* the total enzyme amount, 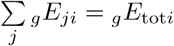, was 101.9 mg.L^−1^, namely the physiological total enzyme concentration. After drawing enzyme concentrations under the different conditions described above (equidistributed/centered on the optima for different *c*_v_ values), the concentration _*g*_*E*_*ji*_ of enzyme *j* in individual *i* was computed as:

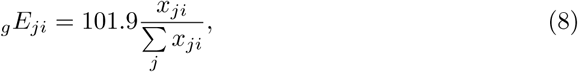

where *x_ji_* was the result of the draw. Thus the total enzyme concentration was 101.9 mg.L^−1^ for every individual. It is worth noting that the constraint on *E*_tot_ changes the *c*_v_’s in an inverse relation to enzyme concentrations: the *c*_v_’s of the most (resp. the less) abundant enzymes will decrease (resp. increase) under constraint (see S1 Appendix).

For every simulation condition, we drew two series of 10 000 parents which were “crossed” to get 10 000 hybrids. Parent and hybrid fluxes were computed according to Eq 6, assuming additivity of enzyme concentrations.

##### Computer simulations of the glycolysis and fermentation network in yeast

We modeled the glycolysis and fermentation network in yeast using a simplified topology in which there was one input flux, the rate of glucose consumption, and two output fluxes, the rate of glycerol production and the rate of transformation of pyruvate into acetaldehyde (Fig 3). The system of ordinary differential equations we used was derived from Conant & Wolfe’s model [80] obtained from the Biomodels database (http://www.ebi.ac.uk/biomodels-main/). This model is a slightly different version of a previously published model [81]. We modified this model in order to increase the range of enzyme concentrations that can lead to a steady state. First, we excluded the reactions producing trehalose and glycogen because, due to their constant rates, they caused incompatibility when the entry of glucose decreased. Second we suppressed mitochondrial shuttling, which is a composite reaction, because it is not associated with any enzyme. Thus our model included 17 reactions and 22 metabolites, including 6 cofactors (ATP, ADP, AMP, NAD, NADH and Fru-2,6-BP) (Fig 3). For all simulation conditions (see below), we solved the system of ordinary differential equations numerically using solvers (deSolve package) of the Runge-Kutta family [82] implemented within the R software tool (R Development Core Team 2010). We considered that the steady state was reached when metabolite concentrations varied less than 10^−4^ between two successive time steps.

**Fig. 3.**
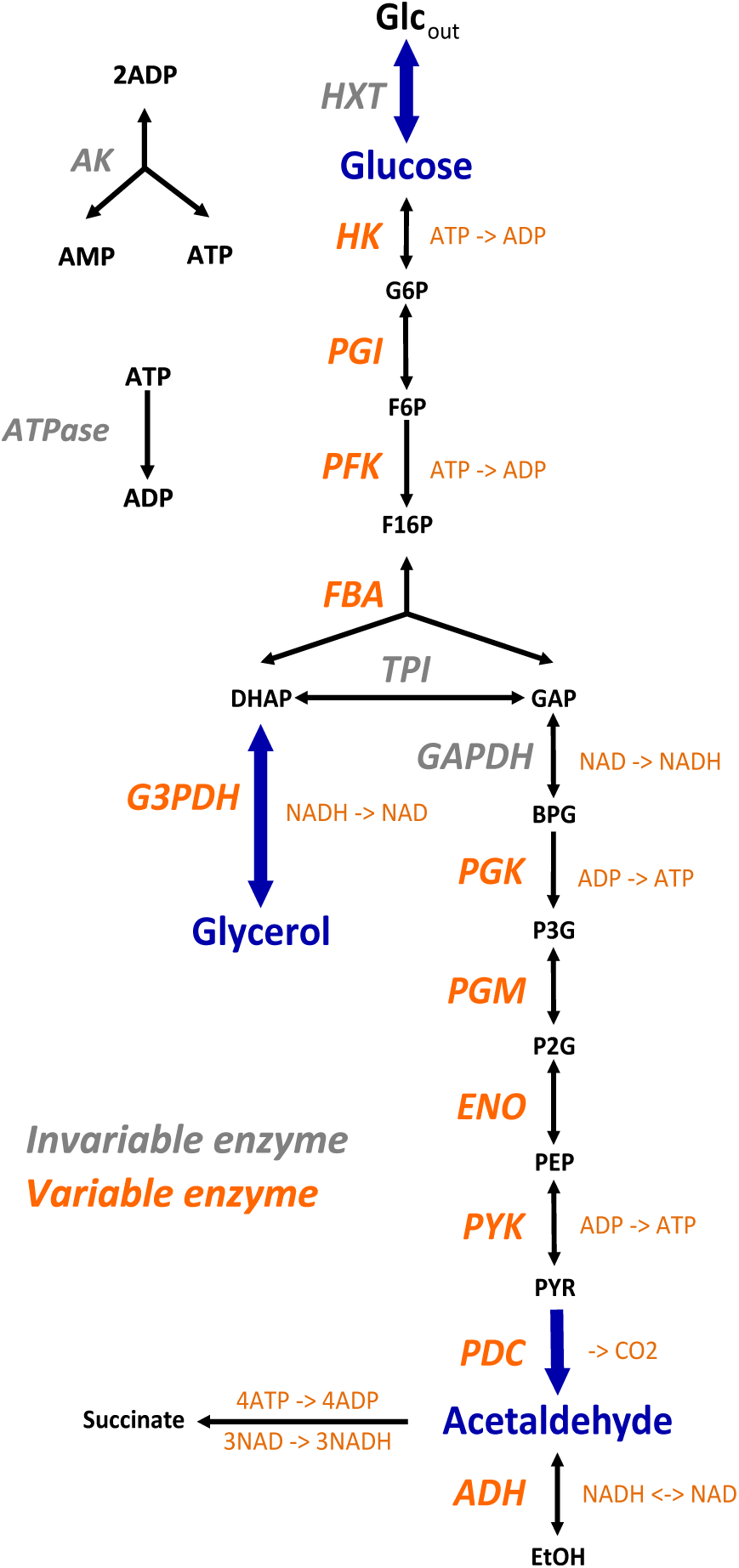
The simplified yeast glycolysis/fermentation network. Variable enzymes are in orange. Blue arrows point to the input flux, glucose, and to the output fluxes, glycerol and acetaldehyde.

Conant & Wolfe’s model uses the maximal enzymatic rate *V*_max_, i.e. there is no explicit expression of enzyme concentration and catalytic constant *k*_cat_. To simulate variation in enzyme concentrations we needed to know the *k*_cat_ values and molecular weights of the enzymes. We found most *k*_cat_ values in the Brenda database (http://www.brenda-enzymes.info/). When they were not known in yeast, we used data from other organisms. We found *k*_cat_ values for 11 enzymes (S3 Table: HK, hexokinase; PGI, phosphoglucose isomerase; PFK, phosphofructokinase (E.C. 2.7.1.11); FBA, fructose-1,6-bisphosphate aldolase; PGK, 3-Phosphoglycerate kinase; PGM, phosphoglycerate mutase; ENO, enolase; PYK, pyruvate kinase; PDC, pyruvate decarboxylase; ADH, alcohol dehydrogenase and G3*P*DH, glycerol 3-phosphate dehydrogenase. For these enzymes, we determined the reference concentrations *E*_ref_ of the model from the equation:

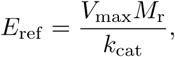

where *M*_r_ is the molecular weight of the enzyme (S3 Table).

As expected, the flux response to variation in concentration of each enzyme, with the other 10 concentrations being maintained at their reference values, displayed saturation curves in most cases. However, for certain enzymes the reference concentration was far or very far along the plateau, so that varying the concentration was almost without effect on the fluxes. Therefore we tried to arbitrarily decrease the reference values of these enzymes, but as a consequence the rate of successful simulations decreased dramatically. In the end, we solved this problem by decreasing the 11 reference concentrations by the same factor, which was empirically fixed at 5. Thus we simulated the genetic variability of parental enzyme concentrations by drawing values from gamma distributions with parameters 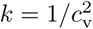 and 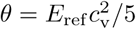. Variances were set to have seven *c*_v_’s ranging from 0.1 to 0.7. With higher *c*_v_’s the steady state was frequently not reached, and *c*_v_’s smaller than 0.1 produced results similar to those obtained with *c*_v_ = 0.1. For each *c*_v_ value, we drew 10 000 pairs of parents. The enzyme concentrations of their hybrids were computed assuming additive inheritance. We considered two situations, free *E*_tot_ and fixed *E*_tot_. For free *E*_tot_, we used the concentration values drawn as described above. For fixed *E*_tot_, the concentrations were scaled to make their sum equal to the sum of the reference enzyme concentrations (31833.97 mg), as previously explained (Eq 8): 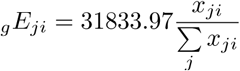. We checked that the input flux (glucose) was equal to the sum of the output fluxes (glycerol and acetaldehyde), indicating that there was no “leak” in the system.

For each parent-hybrid triplet we computed the fluxes of glucose, glycerol and acetaldehyde at the steady state. There were two possible sources of bias on the *a posteriori c*_v_’s. First, if one member of the triplet did not reach a steady state, the triplet was excluded. Thus, to get 10 000 simulations, *≈* 10 060 to *≈* 36 200 simulations were necessary, depending on the *c*_v_ and the presence of constraint on *E*_tot_. Second, even if a steady state was reached, the glucose flux could be null, and we obviously eliminated the corresponding triplets. An analysis of the resulting biases is presented in S2 Appendix. Overall, the differences between *a priori* and *a posteriori c*_v_ values were insignificant or moderate when initial *c*_v_’s were small to medium, and were significant only for certain enzymes with high initial *c*_v_’s. In any case, these differences maintained the wide range of *c*_v_’s, making it possible to study the effects of variability in enzyme concentrations on heterosis.

##### Flux differences and enzymatic distances between parents

The relationship between parental flux values and heterosis was analyzed using the absolute value of the flux difference between parents *i* and *i*′:

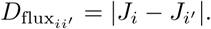

The relationship between parental enzyme concentrations and heterosis was analyzed using the weighted normalized Euclidean distance between parents *i* and *i*′:

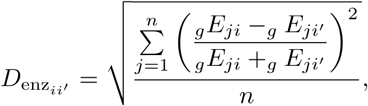

where *n* is the number of enzymes. With normalization, we have 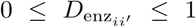.

## Results

In order to analyze the way the genotype-phenotype (GP) relationship shapes heterosis, we used theoretical developments, *in vitro* genetics and computer simulations of the glycolytic/fermentation flux.

### A geometrical approach: the concavity of the GP relationship results in heterosis

As an archetype of the concave GP relationship, we first chose a hyperbolic function relating the flux *J_i_* of individual *i* to enzyme concentrations in a *n*-enzyme metabolic system:

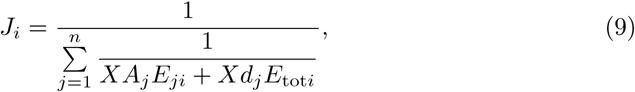

where *X* is a positive constant, *A*_*j*_ and *d*_*j*_ are systemic positive parameters accounting for the kinetic behavior of enzyme *E*_*j*_ in the network, and *E_ji_* and *E*_tot_*i* are respectively the concentration of enzyme *E*_*j*_ and the total enzyme concentration in individual *i* [13, 71, 76]. Because we assumed in this study that there was additivity of enzyme concentrations, the flux of the hybrid of parents P1 and P2, with fluxes *J*_1_ and *J*_2_ respectively, was written:

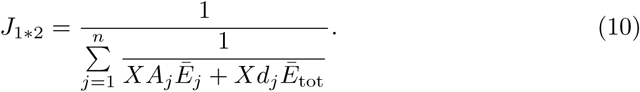

where 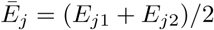 and 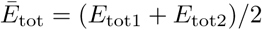. Since the relationship between enzyme concentrations and flux is a rectangular multivariate hyperbolic function, the concavity of the surface results in positive heterosis for the flux 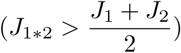, i.e. either positive Mid-Parent Heterosis (+MPH) or Best-Parent Heterosis (BPH) (see S1 Fig for definitions of the types of heterosis). This has been be shown analytically (see Theoretical background), and is illustrated geometrically on the two-dimensional hyperbolic flux response surface obtained with two variable enzymes (Fig 4A). Interestingly, it is possible to show that there is BPH, i.e. *J*_1*∗*2_ *> J*_2_ (for *J*_2_ *> J*_1_), whenever:

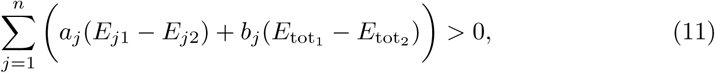

where *a*_*j*_ and *b*_*j*_ are positive terms (see Theoretical background). This equation predicts that heterosis will be observed only if a sufficient number of enzymes have a lower concentration in the parent displaying the highest flux than in the other parent, making (*E*_*j*1_ *− E_j_*_2_) positive, and this effect is strengthened if *E*_tot_ is smaller in the parent that has the highest flux. These conditions are more likely to arise when parental fluxes are similar (Fig 4A). In other words, a condition for observing BPH is that parents display contrasted distributions of enzyme concentrations, i.e. with complementary “high” and “low” alleles.

**Fig. 4.**
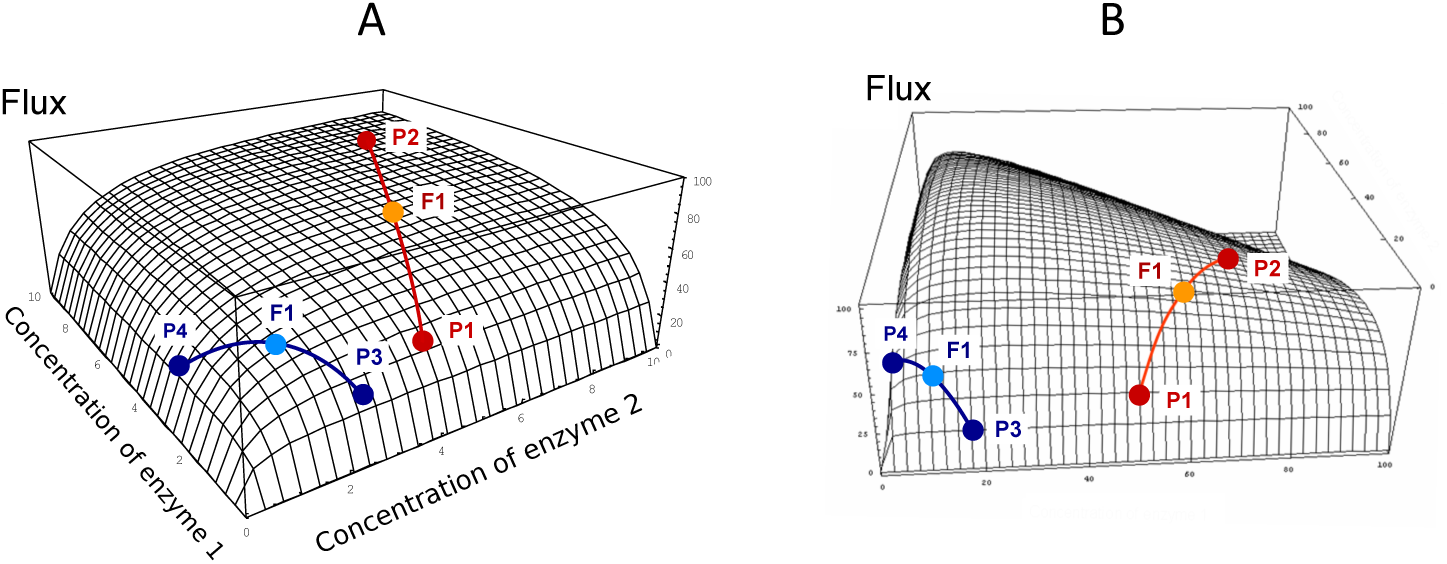
Flux response upon variation of two enzymes in a hyperbolic GP relationship. The two enzymes have the same arbitrary values of their kinetic parameters. Red and blue curves show two types of crosses between parents P1 and P2, or P3 and P4, respectively (from [71]). The positions of hybrids F1 (light blue and orange points) were determined assuming additivity of enzyme concentrations. A: In the “red” cross, where parent P2 has a flux close to the maximum (high concentration of both enzymes), there is positive mid-parent heterosis (+MPH) for the flux. In the “blue” cross, where parents have low flux values due to low concentrations of enzyme 1 (parent P3) or enzyme 2 (parent P4), the hybrid displays best-parent heterosis (BPH). B: If there is a constraint on the total enzyme amount, the “red” cross also displays BPH because the concavity of the surface is increased.

In the case of constrained resources, i.e. if the total enzyme amount allocated to the system is limited, which is biologically realistic, the flux-enzyme relationship is no longer a multivariate hyperbolic function but exhibits a hump-shaped response surface because enzyme concentrations are negatively correlated (Fig 4B). Increasing the concentration of certain enzymes leads to decreasing the concentration of other enzymes [77, 83, 84]. The constraint on *E*_tot_ increases the concavity of the surface, which reinforces the incidence of heterosis (see Theoretical background). If *E*_tot_ is the same in the two parents, the condition for BPH is simply written as:

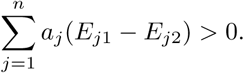

Note that when *E*_tot_ is constant, BPH cannot be observed if a parent has optimal enzyme concentrations, because no offspring can exceed the maximal flux value.

### *In vitro* and *in silico* heterosis in a four-enzyme pathway

In order to assess these predictions, we performed *in vitro* genetic experiments. We reconstructed the first part of glycolysis (Fig 2), and created 61 “hybrid” tubes by mixing 1:1 the content of “parental” tubes that differed in their distribution of the concentrations of four enzymes, PGI, PFK, FBA and TPI, with total enzyme concentration being constant (see Materials and methods).

Inheritance was clearly biased towards high hybrid values (Fig 5). Among the 61 hybrids, 27 (*≈* 44.3 %) exhibited a significantly higher flux value than the mid-parental value: seven (*≈* 11.5 %) displayed BPH and 20 (*≈* 32.8 %) displayed +MPH (*p*-values *<* 0.05). The maximum BPH value, quantified with the index 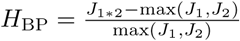, was *H*_BP_ = 0.37, i.e. the hybrid displayed a flux that was 37 % higher than the best parental flux. By contrast, only eight hybrids had a flux value that was significantly lower than their mid-parental value (negative Mid-Parent Heterosis, or –MPH), with no case found of significant Worst-Parent Heterosis (WPH) (Fig 5 and S1 Table).

**Fig. 5.**
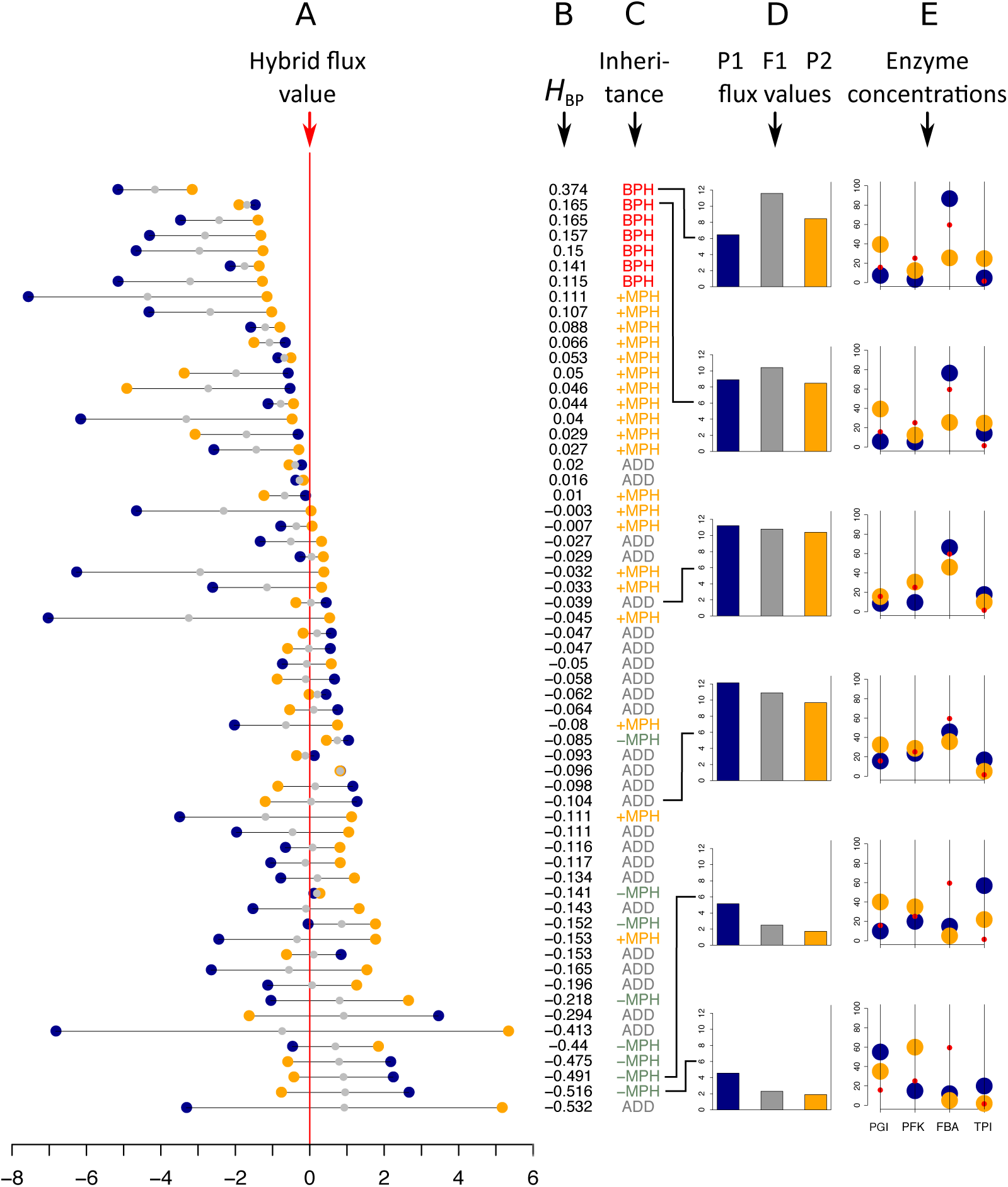
*In vitro* heterosis. Hybrids were created by mixing 1:1 the content of parental tubes. A: Parental flux values (blue and orange points) relative to the hybrid flux value used as a reference (vertical red line) for the 61 crosses (in *µ*M.s^−1^). gray points indicate the mid-parental values. Crosses were ranked by decreasing *H*_BP_ values, the index of best parent heterosis. B: *H*_BP_ values. C: Inheritance: BPH, Best-Parent Heterosis; +MPH, positive Mid-Parent Heterosis; ADD, additivity; –MPH, negative Mid-Parent Heterosis. D and E: Parental (blue and orange bars) and hybrid (gray bars) flux values, alongside the corresponding parental enzyme concentrations for six crosses, identified by broken lines, displaying different types of inheritance. From top to bottom: the two crosses with the highest significant BPH values, the two crosses that are the closest to additivity and the two crosses with the smallest significant –MPH values. Red points correspond to the optimal enzyme concentrations.

Examining enzyme concentrations in the parents confirmed that high values of BPH were likely to occur when parental distributions were contrasted and when neither parent had enzyme concentrations close to the optimal distribution (Fig 5D and E). However, as shown in Fig 6A, this relationship is not straightforward, with large distances being necessary but not sufficient to get high *H*_BP_ values, which resulted in a non-significant correlation between *H*_BP_ and *D*_enz_, the Euclidean distance between parental enzyme concentrations.

**Fig. 6.**
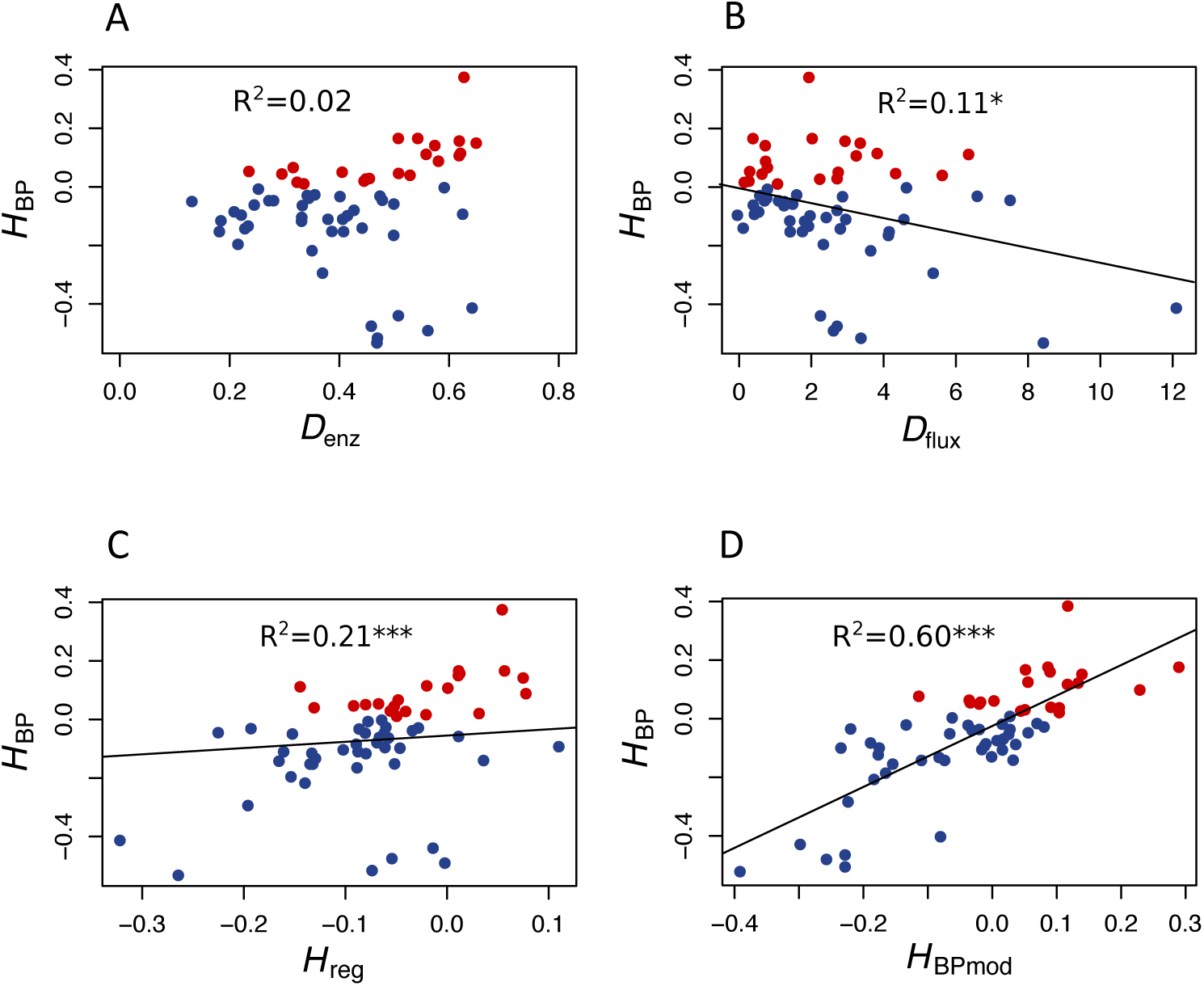
*In vitro* heterosis predictors. A: Relationship between *H*_BP_ and *D*_enz_, the Euclidean distance between parents computed from enzyme concentrations (*r* = 0.15, *p <* 0.25). B: Relationship between *H*_BP_ and *D*_flux_, the flux difference between parents (*r* = −0.33, *p <* 0.01). C: Relationship between *H*_BP_ and *H*_reg_, the heterosis value computed from the equation of the multiple linear regression performed with *D*_enz_ and *D*_flux_ as predictor variables (*r* = 0.46, *p <* 0.001). D: Relationship between *H*_BP_ and *H*_BPmod_, the expected heterosis value computed from the model (see text) (*r* = 0.77*, p <* 2.5.10^−13^). Red points correspond to positive *H*_BP_, i.e. to BPH.

To test the prediction that a cross between parents with similar fluxes is more likely to give heterosis than a cross between parents with contrasted fluxes, we computed the correlation coefficient between *H*_BP_ and *D*_flux_, the difference between parental fluxes (*| J*_1_ *− J*_2_ *|*), and found a highly negative significant value (*r* = −0.33*, p <* 0.01) (Fig 6B).

Interestingly, even though *H*_BP_ and *D*_enz_ were not correlated, the heterosis value *H*_reg_, computed from the equation of the multiple linear regression performed with both *D*_enz_ and *D*_flux_ as predictor variables, was better correlated with *H*_BP_ than *D*_flux_ alone: *r* = 0.46 (*p <* 1.8.10^−4^) (Fig 6C).

Finally, in order to know if the instances of heterosis we observed *in vitro* were consistent with our heuristic model of flux-enzyme relationship, we computed from Eq 9 and Eq 10 the theoretical fluxes of the 61 hybrids and their parents, using the values of enzyme concentrations of the parental tubes and the values of parameters *XA_j_* and *Xd_j_* estimated in an independent experiment (see Materials and methods). Then, for each virtual cross, we calculated the heterosis that would be expected from the model, *H*_BPmod_. As shown in Fig 6D, the correlation between *H*_BPmod_ and *H*_BP_ is very highly significant (*r* = 0.77*,p <* 2.5.10^−13^).

As Eq 9 and Eq 10 resulted in quite reliable predictions of flux heterosis, we used them to simulate a large series of crosses. The parental enzyme concentrations were drawn to be centered on their optimum or equidistributed, and their coefficients of variation (*c*_v_) varied from 0.1 to 1.2. In every case we considered two situations, free and fixed *E*_tot_. For every simulation condition, 10 000 crosses were performed and the parental and hybrid values were computed.

The percentage of BPH and the mean of positive *H*_BP_ values 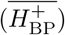 were compared over the range of *c*_v_’s. Fig 7 shows that: (i) in all conditions, the values of these two variables increased, usually to a large extent, when enzyme variability increased; (ii) by far the highest % BPH (about 50 %) was observed when enzyme concentrations were constrained (fixed *E*_tot_) and centred on their optimum (Fig 7A); (iii) 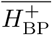 was highly dependent on the *c*_v_ values (varying from *≈* 0 to *≈* 1.4), but was barely affected by the other simulation conditions (Fig 7B).

**Fig. 7.**
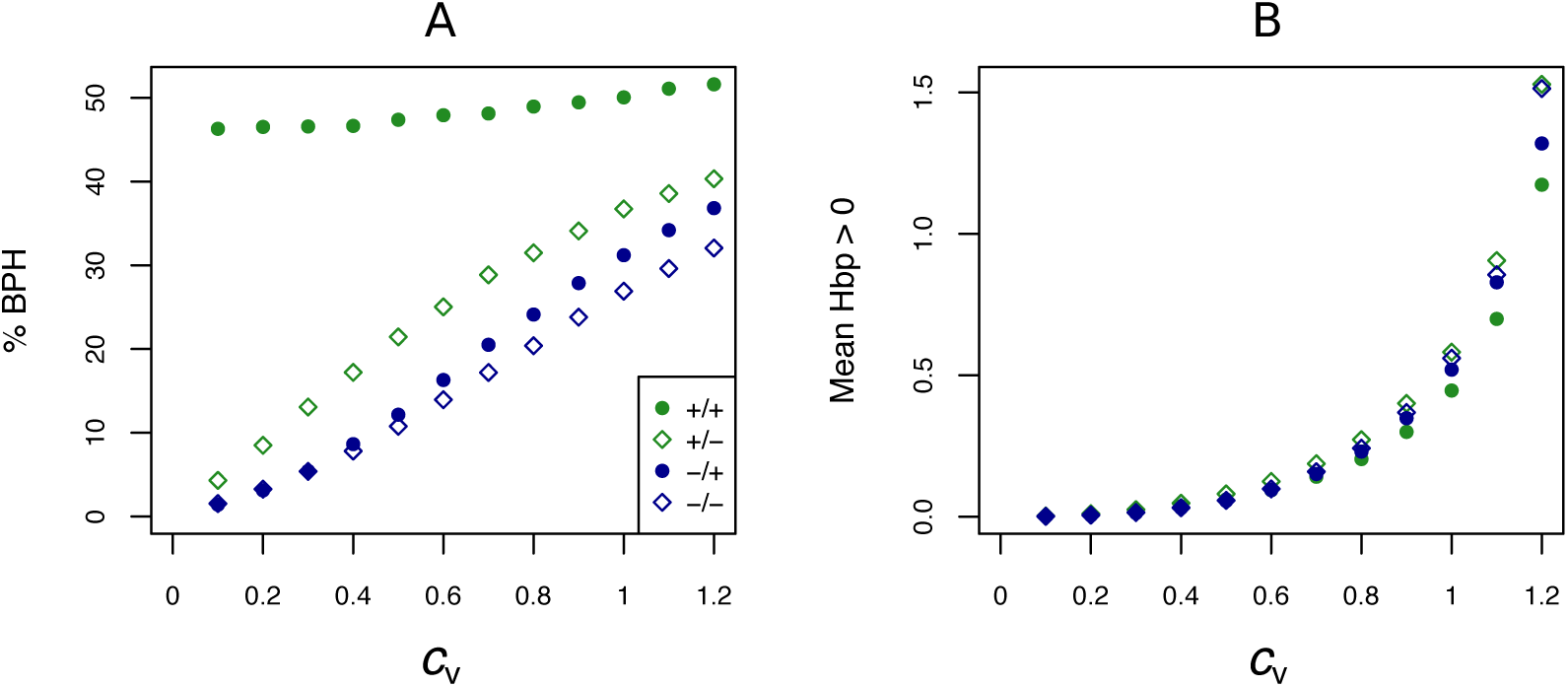
Effect of the coefficient of variation (*c*_v_) of enzyme concentrations on heterosis. A: Percentage of BPH; B: Mean of the positive *H*_BP_ values 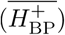 over a range of *c*_v_’s of parental enzyme concentrations, from 0.1 to 1.2. −/−: equidistributed mean enzyme concentrations and free *E*_tot_, −/+: equidistributed and fixed *E*_tot_, +/–: mean enzyme concentrations centered on their optimum and free *E*_tot_, +/+: optimum centered means and fixed *E*_tot_.

The typical influence of *D*_flux_ on *H*_BP_ is illustrated Fig 8 for *c*_v_ = 0.6. BPH was observed when the parental fluxes were close to each other, as the hybrids displaying BPH were distributed around the diagonal of the parental flux space. This pattern is consistent with the negative relationship between *D*_flux_ and *H*_BP_ (Fig 9A). The highest *D*_flux_ never resulted in BPH, while small *D*_flux_ could result in high *H*_BP_ values. The weakest correlation between *D*_flux_ and *H*_BP_ was observed when both enzyme concentrations were centered on their optimum and *E*_tot_ was fixed (*r* = −0.35 instead of *r > −*0.66 in other conditions).

**Fig. 8.**
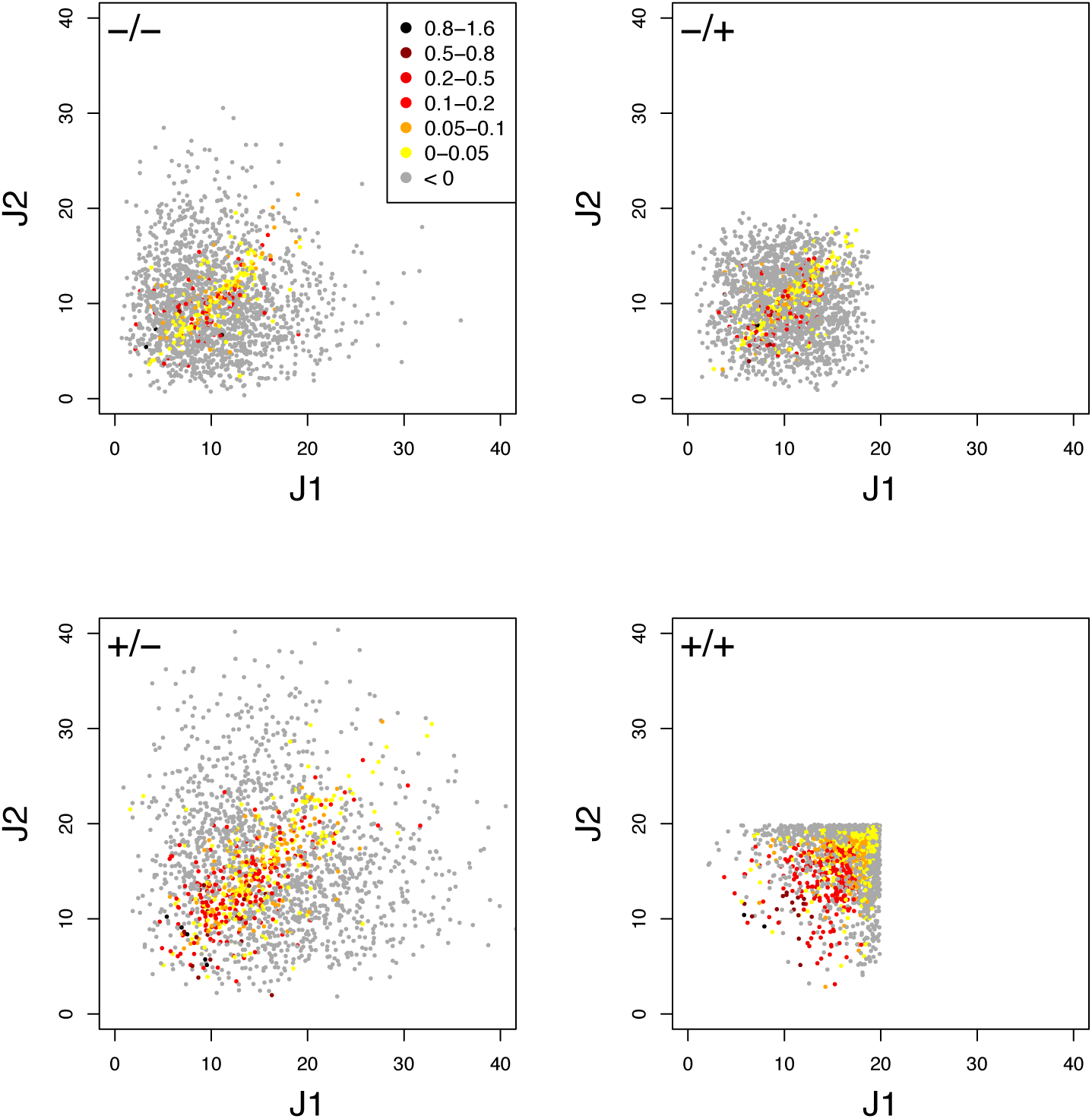
Relationship between parental fluxes *J*_1_ and *J*_2_ and hybrid fluxes under four simulation conditions. Each point corresponds to a hybrid, with warm colors when there is BPH (*H*_BP_ *>* 0). +/+, +/−, −/+ and −/− have the same meaning as in Fig 7. Note that when *E*_tot_ is fixed, the flux value is limited. (*c*_v_ = 0.6.)

The graphs showing the relationship between *D*_enz_ and *H*_BP_ revealed a typical triangular relationship, regardless of the simulation conditions: BPH was never observed with the smallest distances, and large distances were necessary but not sufficient to get high *H*_BP_ values (Fig 9B).

**Fig. 9.**
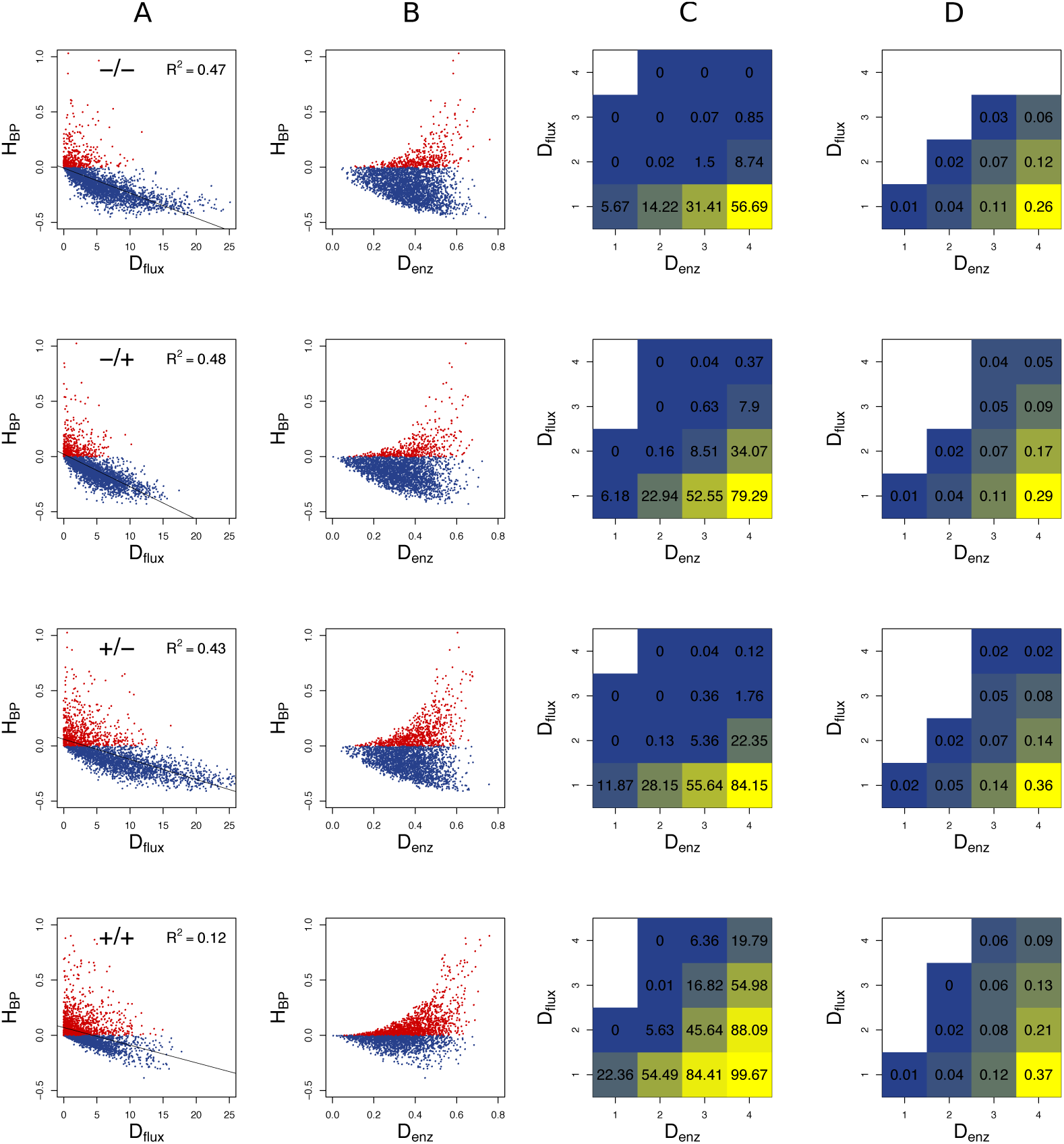
Heterosis predictors. Relationship between *H*_BP_ and *D*_flux_ (A), and between *H*_BP_ and *D*_enz_ (B) when *c*_v_ = 0.6. Red points: positive *H*_BP_ values. C: Square grid of percentages of BPH as a function of *D*_enz_ (x-axis) and *D*_flux_ (y-axis). Colors range from blue (minimum % BPH) to yellow (maximum % BPH). Empty squares are white. D: Same as C with 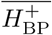 values. The additional white squares correspond to cases where there was no BPH (0 in C). −/−, −/+, +/− and +/+ have the same meaning as in Fig 7.

To assess the joint influence of *D*_flux_ and *D*_enz_ on heterosis, we divided the crosses into 16 classes according to a 4 × 4 grid of *D*_flux_ and *D*_enz_ values, and computed for each class the % BPH and 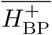. In all conditions, the highest values of these two variables were observed with both small *D*_flux_ and large *D*_enz_ (Fig 9C and D). As expected, constraint on *E*_tot_ resulted in more heterosis, with almost 100 % BPH when: (i) enzyme concentrations were centered on their optimum; (ii) *D*_enz_ values were maximal; (iii) *D*_flux_ values were minimal (Fig 9C). In these conditions, 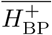 increased to 0.37, i.e. hybrids had on average a flux that was 37 % higher than the best parental flux (Fig 9D).

### *In silico* heterosis in the yeast glycolysis/fermentation network

To simulate heterosis in a larger system, we modeled the glycolysis and fermentation network in yeast using a simplified topology in which there was one input flux, the rate of glucose consumption, and two output fluxes, the production rates of glycerol and acetaldehyde (Fig 3). We used a system of differential equations derived from [80] and varied the concentration of 11 enzymes.

We first examined how varying the concentration of one enzyme at a time, the other concentrations being maintained at their reference values (see Material and methods), affected flux variation. When *E*_tot_ was free, the enzyme-flux relationships were concave with a horizontal asymptote in most cases, although a slight convexity was visible in two instances (S2 Fig): (i) sigmoidal curves were observed upon variation of HK, an allosteric enzyme that is by far the most abundant in the system (S3 Table); (ii) due to the balance between the acetaldehyde and glycerol branches, increasing the concentration of an enzyme in one branch convexly decreases the flux of the other branch (except for enzyme at low concentrations). The decrease of the glucose flux upon G*3*PDH increase is likely to be due to the related reduction of the acetaldehyde flux, which dampens ATP production and in turn reduces the glucose flux.

When *E*_tot_ was fixed, these observations remain qualitatively valid, however we no longer found horizontal asymptotes: due to this constraint, flux values decreased when enzyme concentrations were high (S3 Fig). This effect was marked for the most abundant enzymes (particularly HK and PYK) but was imperceptible for less abundant ones (e.g. PFK). Note that in all situations the three fluxes are null when enzyme concentrations are null, even for enzymes downstream of the bifurcation. Cofactor availability probably explains this observation.

We then simulated 10 000 crosses between parents that differed in their concentrations of the 11 variable enzymes. Parental concentrations were drawn from gamma distributions, the means of which were the reference values. Variances were chosen within the range of seven *c*_v_’s, from 0.1 to 0.7, with free or fixed *E*_tot_. Hybrid fluxes and inheritance were computed for each cross (see Materials and methods).

Unlike the previous four-enzyme model, where the enzyme-flux relationship was concave by design and could only produce BPH or positive MPH, we observed the four possible types of inheritance, BPH, +MPH, –MPH and WPH. For the glucose and acetaldehyde fluxes, inheritance was very biased towards positive heterosis whatever the *c*_v_, since the sum of the percentages of +MPH and BPH varied from 69.4 % to 96.9 %, and there was no or very few cases of WPH. The highest percentage of WPH, observed for acetaldehyde when *c*_v_ = 0.7 with free *E*_tot_, was only 0.09 %, with a very small mean of the negative *H*_WP_ values 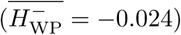 (Tables 1 and 2 and Fig 10). The one-step glycerol branch, which is convexly related to the six enzymes of the acetaldehyde branch, had higher percentages of negative heterosis (from *≈* 65 % when *c*_v_=0.1 to *≈* 43 % when *c*_v_ = 0.7 with fixed *E*_tot_), but WPH remained quite low: 1.23 % and 3.02 % for free and fixed *E*_tot_, respectively, with 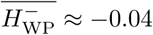 in both cases (*c*_v_ = 0.7). For all fluxes, constraint on *E*_tot_ decreased the percentage of –MPH in favor of BPH and/or +MPH, depending on the *c*_v_ value (Tables 1 and 2 and Fig 10).

**Table 1.** Percentage of occurrence of the four types of inheritance for the glucose, glycerol and acetaldehyde fluxes over the range of *c*_*v*_ values when *E*_tot_ is free.

| $c_v$ | | 0.1 | 0.2 | 0.3 | 0.4 | 0.5 | 0.6 | 0.7 |
| --- | --- | --- | --- | --- | --- | --- | --- | --- |
| Glucose | BPH | 3.76 | 13.80 | 19.77 | 23.25 | 25.44 | 29.97 | 35.14 |
|  | +MPH | 65.67 | 59.44 | 55.89 | 56.12 | 56.04 | 54.09 | 50.35 |
|  | -MPH | 30.57 | 26.76 | 24.34 | 20.64 | 18.51 | 15.94 | 14.52 |
|  | WPH | 0.00 | 0.00 | 0.00 | 0.00 | 0.01 | 0.04 | 0.00 |
| Glycerol | BPH | 2.92 | 10.84 | 14.92 | 16.94 | 16.99 | 18.39 | 18.85 |
|  | +MPH | 31.77 | 33.61 | 33.49 | 35.12 | 38.38 | 37.41 | 38.27 |
|  | -MPH | 64.83 | 54.85 | 50.66 | 47.01 | 43.40 | 43.14 | 41.65 |
|  | WPH | 0.48 | 0.70 | 0.93 | 0.93 | 1.23 | 1.06 | 1.23 |
| Acetaldehyde | BPH | 4.05 | 14.63 | 21.23 | 24.88 | 28.05 | 32.71 | 37.76 |
|  | +MPH | 68.10 | 60.42 | 56.15 | 55.79 | 53.92 | 52.49 | 48.45 |
|  | -MPH | 27.86 | 24.93 | 22.62 | 19.32 | 18.03 | 14.80 | 13.69 |
|  | WPH | 0.00 | 0.01 | 0.00 | 0.01 | 0.00 | 0.00 | 0.09 |

**Table 2.** Percentage of occurrence of the four types of inheritance for the glucose, glycerol and acetaldehyde fluxes over the range of *c*_*v*_ values when *E*_tot_ is fixed.

| $c_v$ | | 0.1 | 0.2 | 0.3 | 0.4 | 0.5 | 0.6 | 0.7 |
| --- | --- | --- | --- | --- | --- | --- | --- | --- |
| Glucose | BPH | 11.96 | 25.86 | 32.89 | 35.58 | 37.57 | 39.14 | 42.46 |
|  | +MPH | 84.92 | 70.44 | 62.00 | 58.60 | 54.89 | 52.90 | 49.29 |
|  | −MPH | 3.12 | 3.70 | 5.11 | 5.82 | 7.53 | 7.95 | 8.25 |
|  | WPH | 0.00 | 0.00 | 0.00 | 0.00 | 0.01 | 0.01 | 0.00 |
| Glycerol | BPH | 3.71 | 13.64 | 18.72 | 20.07 | 21.57 | 23.19 | 24.10 |
|  | +MPH | 31.03 | 30.33 | 31.54 | 33.10 | 34.36 | 34.46 | 35.78 |
|  | −MPH | 64.16 | 53.74 | 46.95 | 43.98 | 41.12 | 39.51 | 37.09 |
|  | WPH | 1.10 | 2.28 | 2.79 | 2.84 | 2.94 | 2.84 | 3.02 |
| Acetaldehyde | BPH | 12.61 | 26.93 | 33.77 | 36.83 | 38.91 | 41.01 | 44.40 |
|  | +MPH | 83.23 | 68.28 | 60.07 | 56.40 | 52.63 | 50.62 | 46.71 |
|  | −MPH | 4.16 | 4.80 | 6.16 | 6.77 | 8.46 | 8.35 | 8.89 |
|  | WPH | 0.00 | 0.00 | 0.00 | 0.00 | 0.00 | 0.01 | 0.00 |

**Fig. 10.**
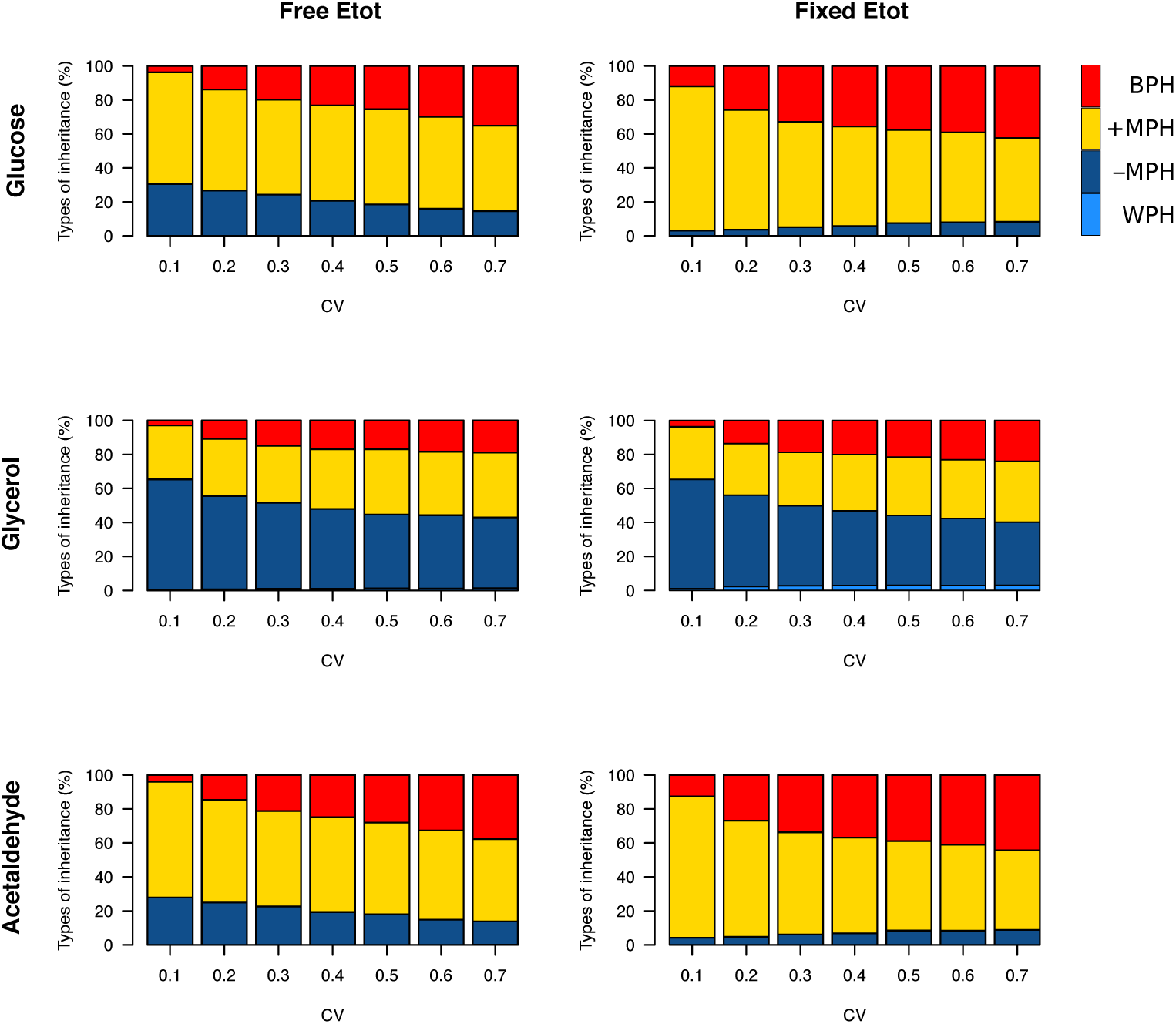
Heterosis for the three fluxes in the glycolytic/fermentation network. Percentages of different types of inheritance over the 0.1–0.7 *c*_v_ range of parental enzyme concentrations, for free and fixed *E*_tot_.

We hypothesized that negative heterosis can occur only when the enzyme-flux relationship is convex, as exemplified for negative dominance (Fig 1A). To verify this, we ran simulations where HK concentration was maintained at its reference value and/or the glycerol branch was deleted. When both sources of convexity were removed, no case of negative heterosis was observed (S4 Table and S4 Fig).

As previously, both % BPH and 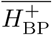 increased as the *c*_v_ increased (Tables 1, 2 and 3, Fig 11A and B). This effect was larger for glucose and acetaldehyde than for glycerol. For instance, for glucose the % BPH varied from 3.76 % to 35.12 % with free *E*_tot_ and from 11.96 % to 42.46 % with fixed *E*_tot_, while for glycerol it varied from 2.92 % to 18.84 % with free *E*_tot_ and from 3.71 % to 24.10 % with fixed *E*_tot_. For the three fluxes, constraint on *E*_tot_ increased the % BPH, as expected, but decreased 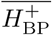. The latter observation is explained by the fact that this constraint decreases the variance of the concentrations, which in turn decreases the flux variance and hence both *H*_BP_ variance and 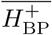 (see S1 Appendix and S2 Appendix).

**Table 3.** Mean of positive *H*_BP_ values 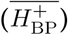 for the glucose, glycerol and acetaldehyde fluxes over the range of *c*_v_ values.

| $c_v$ | | 0.1 | 0.2 | 0.3 | 0.4 | 0.5 | 0.6 | 0.7 |
| --- | --- | --- | --- | --- | --- | --- | --- | --- |
| Free $E_{tot}$ | Glucose | 0.02 | 0.08 | 0.14 | 0.17 | 0.22 | 0.24 | 0.32 |
|  | Glycerol | 0.03 | 0.09 | 0.15 | 0.17 | 0.19 | 0.20 | 0.24 |
|  | Acetaldehyde | 0.02 | 0.09 | 0.14 | 0.18 | 0.23 | 0.28 | 0.38 |
| Fixed $E_{tot}$ | Glucose | 0.01 | 0.04 | 0.08 | 0.12 | 0.16 | 0.21 | 0.25 |
|  | Glycerol | 0.02 | 0.05 | 0.09 | 0.10 | 0.13 | 0.14 | 0.17 |
|  | Acetaldehyde | 0.01 | 0.04 | 0.09 | 0.13 | 0.18 | 0.24 | 0.29 |

**Fig. 11.**
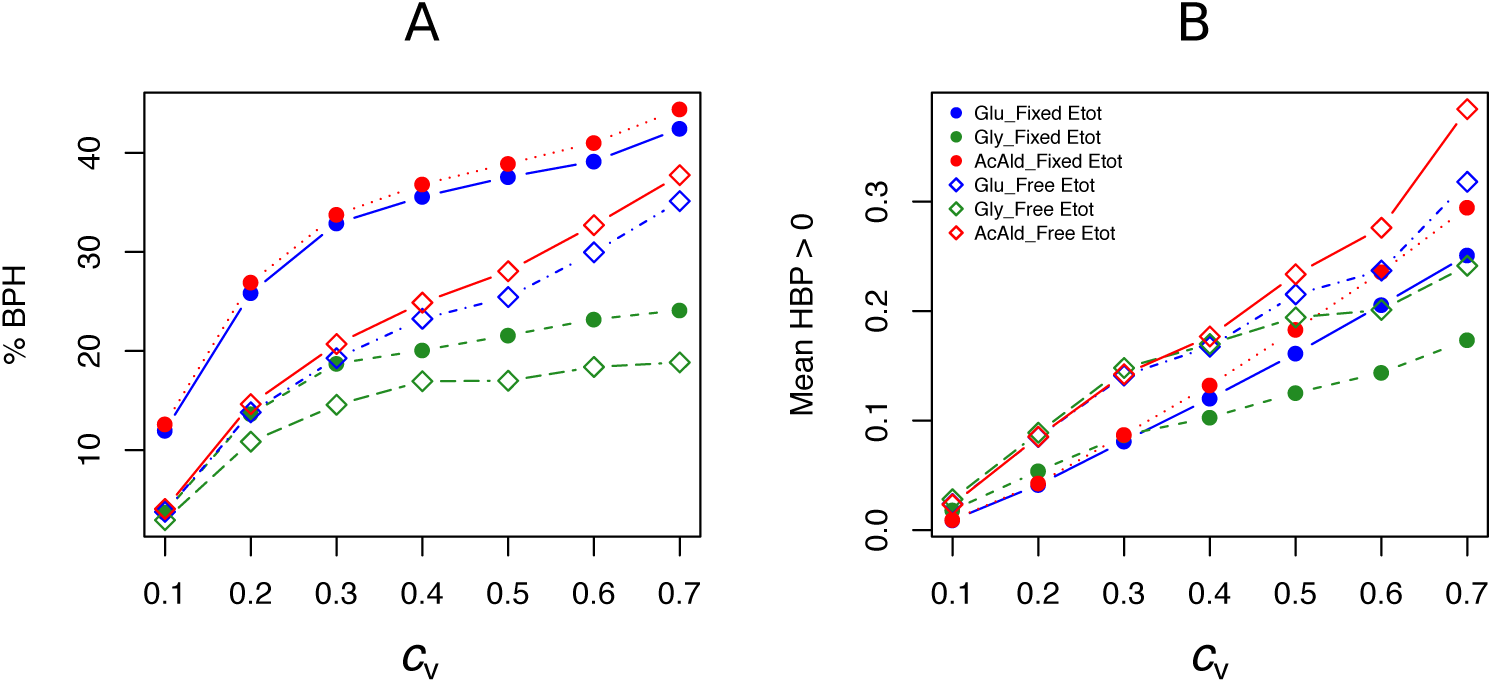
Effect of the coefficient of variation (*c*_v_) of enzyme concentrations on heterosis in the glycolysis/fermentation network. Percentage of BPH (A) and 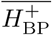 (B) over the *c*_v_ range of parental enzyme concentrations.

Whatever the *c*_v_, BPH was usually observed when parental fluxes were similar, with heterotic hybrids roughly distributed around the diagonal of the parental flux space, and this was valid for the three fluxes (S5 Fig). The correlations between *H*_BP_ and *D*_flux_ were consistent with these observations. They were negative and highly significant, although slightly smaller when *E*_tot_ was fixed (*r* = −0.60*, −*0.69 and −0.61 for glucose, glycerol and acetaldehyde, resp.) than when *E*_tot_ was free (*r* = −0.55*, −*0.59 and −0.54, resp.) (Fig 12A and C).

**Fig. 12.**
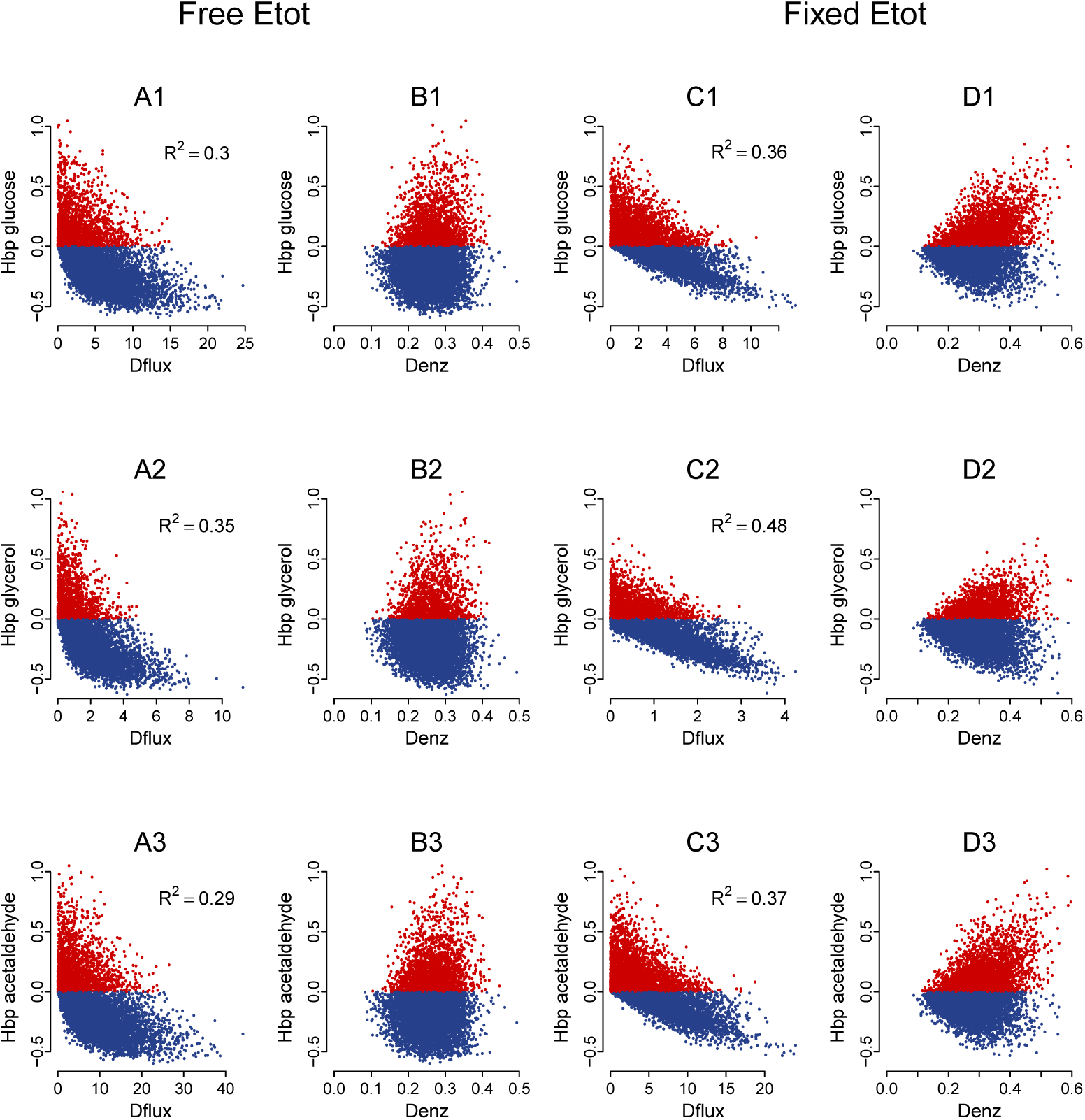
Heterosis predictors in the glycolysis/fermentation network: scatter plots. Relationships between *H*_BP_ and *D*_flux_ (A and C) and between *H*_BP_ and *D*_enz_ (B and D) with free *E*_tot_ (A and B) and fixed *E*_tot_ (C and D) for the glucose flux (A1 to D1), glycerol flux (A2 to D2) and acetaldehyde flux (A3 to D3) (*c*_v_ = 0.4).

The triangular relationship between *H*_BP_ and *D*_enz_ was slightly less apparent with free *E*_tot_ than in the four-enzyme model, but was obvious with fixed *E*_tot_. Again high *H*_BP_ values were never observed with small distances, and the highest *H*_BP_ values could only be observed with medium or large distances (Fig 12B and D).

As previously, we assessed the joint influence of *D*_flux_ and *D*_enz_ on % BPH and 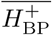 using 4 × 4 grids (Fig 13, for *c*_v_ = 0.4). Results were very similar: both variables were positively related to *D*_enz_ and negatively related to *D*_flux_, whatever the flux. High *D*_enz_ associated with small *D*_flux_ could result in about 50 % BPH and 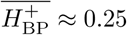 when *E*_tot_ was free and almost 100 % BPH and 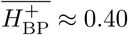 when *E*_tot_ was fixed. The positive effect of the constraint on 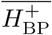 is not in contradiction with the above-mentioned argument: 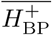 was computed here for each square of the grid, and its distribution was more uneven under constraint. On average 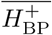 was lower under constraint, but reached higher values when there was both small *D*_flux_ and high *D*_enz_.

**Fig. 13.**
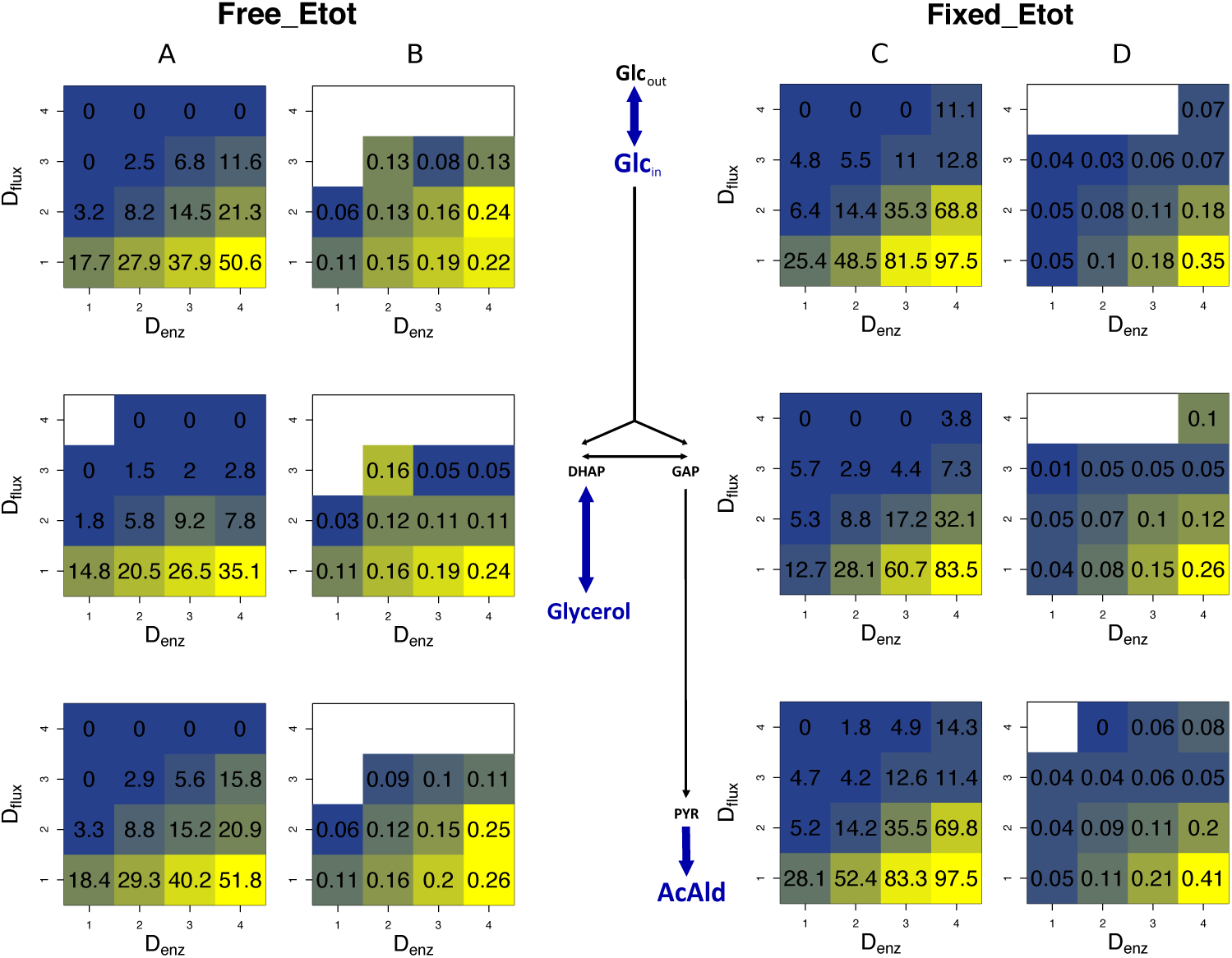
Heterosis predictors in the glycolysis/fermentation network: square grids. A and B: Square grid with % BPH (A) and 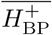 (B) as a function of *D*_enz_ (x-axis) and *D*_flux_ (y-axis) for glucose (1st row), glycerol (2nd row) and acetaldehyde (3rd row) when *E*_tot_ is free. C and D: Same as A and B with fixed *E*_tot_.

S6 Fig shows that contrasted distributions of enzyme concentrations can lead to BPH whereas close distributions favor additivity. Indeed, the cross that is the most heterotic for both glucose and acetaldehyde (*H*_BP_ = 4.23 and *H*_BP_ = 5.55, resp.) had divergent enzyme concentrations (*D*_enz_ = 0.68) (S6 FigA and B); this was also observed for the most heterotic cross for glycerol (*D*_enz_ = 0.66) (S6 FigC and D). These enzymatic distances are close to the highest distance observed (*D*_enz_ = 0.71), which also corresponded to large heterosis for glucose and acetaldhyde (*H*_BP_ = 0.69 and *H*_BP_ = 1.09, resp.) (S6 FigE and F). By contrast, the cross with the minimum enzymatic distance (*D*_enz_ = 0.16) exhibited additivity or near additivity for the three fluxes (*H*_MP_ = 0, *H*_MP_ = 0.02 and *H*_MP_ = −0.01 for glucose, glycerol and acetaldehyde, resp.) (S6 FigG and H).

The geometric models of Fig 14 and S7 Fig show a relationship between *D*_enz_, *D*_flux_ and *H*_BP_ that is fully consistent with all our observations. For a given value of *D*_enz_, *H*_BP_ can be positive or negative depending on the position of the parents in the enzyme concentration space, with quite complex relationships between these two variables. BPH is observed when parents have complementary enzyme concentrations, with the highest *H*_BP_ values observed when *D*_flux_ is small. Of note, *D*_flux_ and *H*_MP_ are roughly positively related, because the reference for this index is the mid-parent and not the best parent (Fig 14E and F). Regarding the constraint on *E*_tot_, comparison of Fig 14A and S8 Fig shows how greater surface curvature increases the incidence of heterosis.

**Fig. 14.**
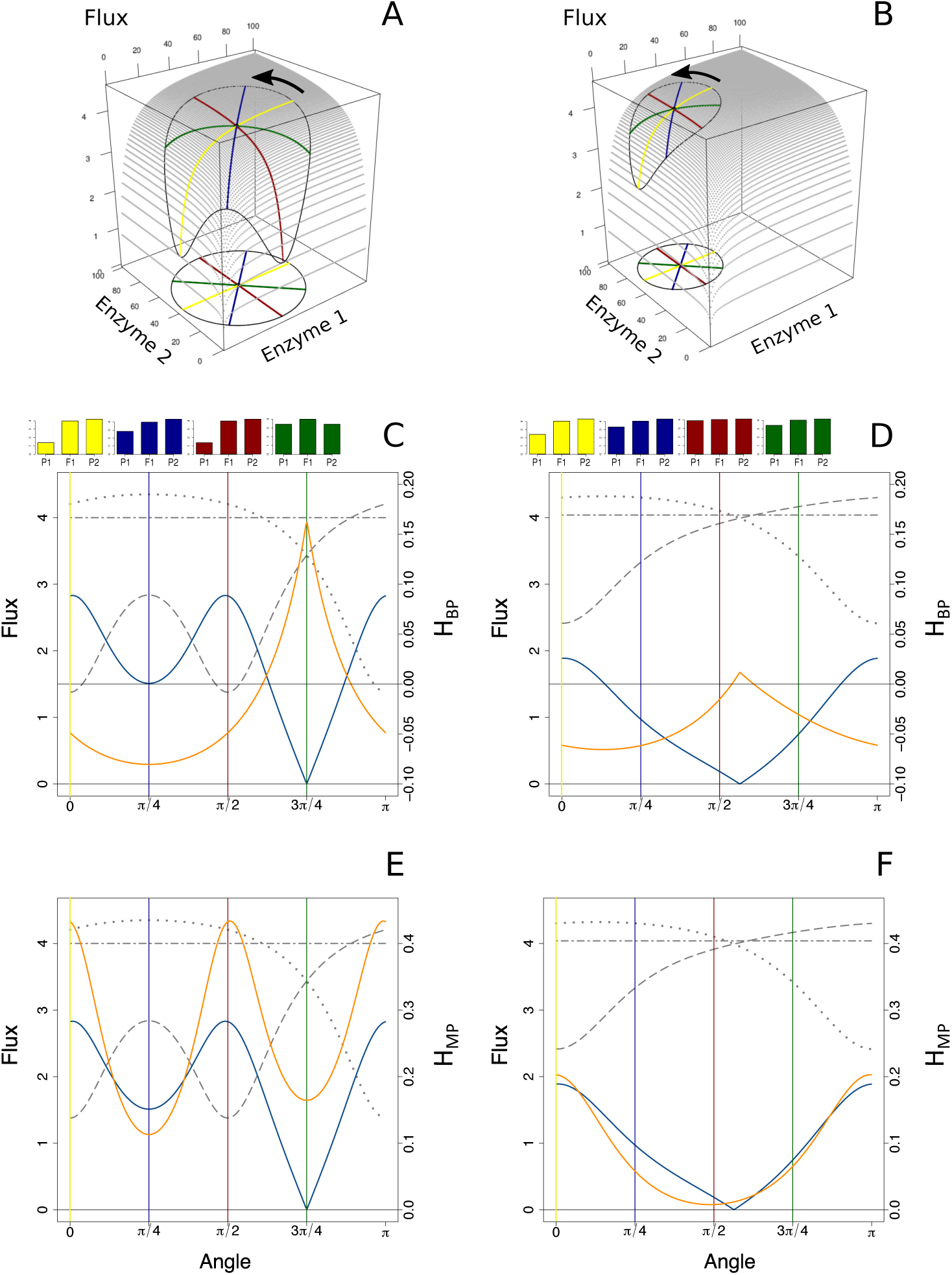
The geometry of heterosis when the enzymatic distance between parents is constant. A: Parental enzyme concentrations are on a circle of diameter *D*_enz_ = 76 centered on the hybrid point, the coordinates of which are *x* = *y* = 40. Four pairs of parents are highlighted by yellow, blue, red and green diameters. The closed curve on the surface shows the flux values corresponding to the circle of concentrations. The equation of the flux surface is 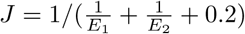, in arbitrary units. B: Same as A, except that *D*_enz_ = 50 and hybrid coordinates are *x* = 30 and *y* = 70. C: Variation of parental fluxes *J*_1_ and *J*_2_ (dotted and long-dashed gray curves), of *D*_flux_ (blue curve) and of *H*_BP_ (orange curve) over the rotation of a segment joining the parents from *α* = 0 (parallel to the x-axis, yellow case) to *α* = *π* (reversed positions of the parents). The two-dashed horizontal gray line corresponds to the constant hybrid flux value. The yellow, blue, red and green vertical lines correspond to the pairs of parents of the 3D vignette in A, with barplots showing corresponding heterosis. D: Same representation as in C, for the case shown in B. E and F: Same as in C and D, with *H*_MP_ instead of *H*_BP_.

## Discussion

### The so-called mystery of heterosis

Nearly all papers on heterosis mention in their introduction that is it an elusive, enigmatic or even magical phenomenon, whose molecular (or physiological, or biochemical) bases are not known or not well understood, and which would be the “lasting mystery in biology”. Yet, this “fascination with the idea that standard genetic models are not sufficient to explain heterosis” [85] is quite surprising given the literature on this phenomenon.

First, regardless of any genetic or molecular basis, it was recognized early on [86,87] that multiplicative traits, i.e. traits determined as the product of two or more components, display commonly predictable levels of heterosis. For instance, heterosis for plant height in bean is quite well explained by multiplying internode length by internode number [88]; similarly, yield heterosis in barley is predicted by the product of ears/plant, kernels/ear and average kernel weight [89], etc. It is worth noting that this cause of heterosis does not even require non-additive inheritance of the components of the trait (reviewed in [90]).

Second, heterosis has been explained at the genetic and/or molecular level for various traits in various species. Paradoxically, the best understood cases of heterosis are certainly the least common, since they involve only one locus, which makes them easier to study. Here, heterosis is due to overdominance, i.e. the inherent superiority of the heterozygote over both homozygotes [91, 92]. Relative to the innumerable examples of complete or partial dominance, well-attested overdominance remains quite rare, with probably less than twenty cases described over more than one century of genetics. The bases of overdominance are diverse: (i) Pleiotropy. The individual traits controlled by the gene harbor only dominance, but the direction of dominance is reversed between the traits, which results in increased viability or fitness of heterozygotes compared to the parental homozygotes [93–95]; (ii) Dosage effects. When there is an optimal expression level of the gene controlling the trait, the hybrid will be heterotic whenever its expression level will be closer to the optimum than that of its parents [96–98]; (iii) Formation of favorable oligomers in the hybrids [99, 100].

Understanding heterosis when several – possibly a wealth of – polymorphic genes control the trait is challenging. However this has been done when a limited number of genes are involved. As early as 1910, an example of heterosis accounted for by complementary dominance at two loci *L* (controlling internode length) and *T* (controlling stem thickness) was found in garden pea [101], in line with Davenport’s model [102], which stated that heterosis was due to recessive unfavorable alleles being masked at loci in repulsion. Since then, similar examples have been described (e.g. [103]). It is worth noting that many alleged overdominant QTLs (Quantitative Trait Loci) could actually correspond to pseudo-overdominance, where two (or more) linked dominant QTLs are in repulsion [104–107]. Epistasis also accounts for heterosis through favorable intergenic interactions created in the hybrids [108]. The textbook case [109], which involves both dominance and epistasis, was recognized very early on [110] and was illustrated in various plants: crossing two individuals with colorless flowers may produce hybrids with colored flowers because the parents each have a mutation inactivating a particular step of the pigment biosynthesis pathway, which disrupts the flux, and the flux is restored in the hybrid because both steps are active (e.g. [111] for anthocyanins in maize). Other cases of heterosis that have been elucidated at the molecular level also combine more than one genetic effect. For instance in yeast, heterosis for growth at high temperature involved both dominance and epistasis at three genes *MKT*1, *END*3, and *RHO*2 [68, 112], and heterosis for fermentation performance proved to be partially explained by dominance and pseudo-overdominance at four genes *VMA*13, *PDR*1, *PMA*1 and *MSB*2 [107]. Similar complex effects were found in the circadian clock genes *CCA*1 and *LHY*, which play a central role in heterosis for chlorophyll and starch content in *A. thaliana* [61]. In tomato, epistasis and overdominance at the *SFT* gene account for yield heterosis [98].

Third, given the pervasiveness of non-additive genetic effects [113, 114], it is the *lack* of heterosis that would be a mystery. Whenever there is dominance, and/or overdominance and/or epistasis at loci governing a quantitative trait, heterosis can arise. Any one of these three genetic effects is necessary, but also sufficient, to generate heterosis. Besides, their respective importance in producing heterosis is variable, and depends on the mating system, the traits under study and the genetic material used, as shown from QTL/association mapping analyses and other approaches. In maize, dominance seems to be the main factor producing heterosis for grain yield and its components, with a modest role of overdominance and possibly epistasis [115–117]. In rice, dominance was first invoked as the main factor for heterosis of grain yield and its various components [57], but subsequently overdominance and additive x additive or dominance x dominance epistasis have been repeatedly found for various yield-related traits [115, 118–123]. In Arabidopsis the main factor was claimed to be epistasis for seven growth-related traits [124], however for biomass the three genetic effects seemed to be involved [125], and a recent study showed the role of dominance and overdominance for flowering date, rosette diameter and rosette biomass [126]. In upland cotton, the three effects contribute to heterosis for yield and yield components [127]. In yeast, the three effects also play a role in heterosis for growth rate measured in five different culture media [128], while overdominance and synergistic epistasis seem mainly involved in growth at high temperature [129]. It is worth noting that in most cases it is not possible to distinguish true overdominance from pseudo-overdominance, even if indirect arguments suggest true overdominance [130].

Despite the multiple cases where the bases of heterosis have been clarified, some authors have been searching for a “unifying principle” [131], such as genome-wide changes in DNA methylation [132,133], small RNA expression and epigenetic regulation [134–136], reduced metabolic cost of protein recycling in hybrids [137], gene dosage effects in macro-molecular complexes [138], enhanced metabolic efficiency due to weak co-aggregation of allozymes [139], organellar complementation [140] and phytohormonal expression [141]. These molecular processes may indeed distinguish heterozygotes from homozygotes, but if they were the general hidden causes of heterosis, correlations between levels of heterosis for different traits would be observed, and this has never been reported so far [142, 143]. In fact, searching for the mechanistic bases of heterosis of a certain trait in a certain cross is equivalent to searching for the bases of its underlying genetic effects, and dominance, epistasis and overdominance have never been considered “mysterious” phenomena.

### Curvature of the genotype-phenotype relationship and heterosis

The “unifying principle” proposed in this paper is not mechanistic. It is a general framework built on the common observation of the concavity of the GP relationship. The two genetic consequences of this non-linearity – dominance and epistasis – have been known for a long time in the context of the metabolic control theory [1, 34]. More recently, modeling gene and metabolic networks have led to the idea that heterosis could also be a systemic property emerging from non-linear processes in the cell [69, 71]. We provide strong arguments in support of this proposition by taking advantage of theoretical developments, and using both experimental approaches and computer simulations.

Experimentally, we performed *in vitro* 61 crosses between “parents” that differ in the concentrations of four enzymes from the upstream part of the glycolysis pathway, and observed inheritance that was biased towards positive heterosis despite additivity of enzyme concentrations in the hybrid tubes: about 44 % of hybrids displayed positive MPH or BPH, whereas only 13 % displayed negative MPH and none WPH. In addition, the *H*_BP_ index could increase up to *≈* 0.37, which means that with enzyme concentrations half-way between the parents, the hybrid had a flux that was 37 % higher than the best parental flux. From a geometrical standpoint, these results indicate that the GP hypersurface in this *in vitro* system is globally concave, accounting for the predominance of positive heterosis, the cases of negative heterosis revealing local unevenness of the surface. We did not explore further the possible causes – metabolic and/or technical – of these irregularities.

Computer simulations based on the previous system and using a hyperbolic GP relationship allowed us to perform a large number of crosses. Enzyme concentrations were drawn, varying means and coefficients of variation, and total enzyme concentrations were either free or fixed. In all conditions only +MPH and BPH were observed, which was expected due to the concavity of the GP relationship. The results of the simulations together with our theoretical developments allowed us to identify the factors favoring heterosis:

A contrast between parental enzyme concentrations. A high positive *H*_BP_ could only be observed when *D*_enz_ was large. Accordingly high *c*_v_’s resulted in the highest % BPH and *H*^+^. However, high *D*_enz_ is necessary but not sufficient to get BPH: the position of the parents in concentration space is also a key factor (Fig 14 and S7 Fig). If a parent contains most of the high alleles and the other most of the low alleles, BPH is unlikely because the hybrid can hardly exceed the best parent, while if the parents are complementary for the high and low alleles, the hybrid is expected to display BPH.
Phenotypic proximity between parents. In the case of strict concavity, BPH is very likely when the parents have close phenotypic values, and the % BPH decreases as the parental fluxes diverge. If a parental value is close to the maximum, MPH alone will be observed (Fig 14).
Constraint on total enzyme concentration allocated to the system. All other things being equal, the % BPH is higher when *E*_tot_ is fixed, due to increased concavity of the flux response surface (Fig 4B).

Combining fixed *E*_tot_, a large enzymatic distance and a small phenotypic difference makes BPH almost inevitable, particularly when mean enzyme concentrations are centered on their optimal values. This is consistent in geometric terms: when these four conditions are met, the hybrids are closer to the “top of the hill” than their parents (S8 Fig).

We performed similar simulations from a branched network based on glycolysis and fermentation in yeast, modeled with a system of differential equations. Three fluxes were considered: glucose input and glycerol and acetaldehyde outputs. Again BPH could only arise when the parents were phenotypically close to each other, but due to some convex enzyme-flux relations in this system, negative heterosis was also observed. For glucose and acetaldehyde fluxes, positive MPH and BPH prevailed to a large extent, since the percentage of positive heterosis varied from 69,5 % to 96.9 %, and for all fluxes and conditions the % BPH increased with the *c*_v_, up to *≈* 44 % when *E*_tot_ was fixed. For glycerol, positive heterosis was predominant only for medium and high *c*_v_’s. In all cases WPH remained very rare. Thus the three fluxes, even though they are pleiotropically related, do not have the same type of inheritance. However, the factors favoring BPH are the same as before for the three fluxes, namely constraint on *E*_tot_ and a large enzymatic distance together with a small phenotypic divergence. Therefore these factors do not depend on a particular flux-enzyme function, but only on the shape of the GP relationship. In addition we see that heterosis is a systemic property which enables a better exploitation of cell resources, since with a distribution of enzyme concentrations closer to the optimum distribution than both parents, a hybrid can “do more with less”, i.e. exceed the best parent with a lower total enzyme concentration. In epistemological terms, it is an *emergent* property, in the sense that the individual properties of the molecular components (transcription/translation factors, mRNAs, proteins/enzymes, metabolites, etc.) cannot alone account for the phenomenon.

In order to avoid confounding effects, in particular in the glycolysis/fermentation system, we did not consider in this study possible non-additive inheritance of enzyme concentrations. Non-additivity of genetic variables has logical consequences: if it is positive, there is a higher occurrence of positive heterosis, while negative non-additivity can result in negative heterosis even in the case of a strictly concave GP relationship [71,74]. In actual fact, transcripts and proteins display mostly additive inheritance [72], and in the case of non-additivity it is usually biased towards positive values [74,133,136,144–147], which will reinforce positive heterosis. For certain gene/protein categories that are found to be non-additive (histone modifications, small RNAs, protein and carbon metabolism, ribosome proteins, photosynthesis, stress response and energy production), a causal link with heterosis has been suggested or evidenced [136,146]. Incidentally, positive non-additivity at the molecular level also reflects concave GP relationships: transcript, protein and metabolite abundance are polygenic traits [148–150] that behave like any quantitative trait and can display non-additive inheritance.

How can our results be applied to whole-cell metabolism? Crowding [151] and energy cost [152,153] create constraints on the total content of cell components. For instance in yeast, Albertin et al. [79] showed that the enzymatic pool allocated to the glycolysis, glycerol, acetate, and ethanol pathways was invariant regardless of the culture medium and strain considered. When some proteins are overexpressed, this constancy can sometimes lead to burden effects, which causes a decline in flux [154], as we found under constrained *E*_tot_. However, due to the huge number of molecular species in a cell, the increase of particular components does not usually impair the whole system. Similarly, situations of convexity within cell networks are expected to be buffered at the scale of the whole flux of matter and energy through the system. Thus, despite global constraints and local convexity, plateaued GP relationships, either S-shaped or more often concave, are commonly observed at integrated levels of cell organization, up to fitness, and most likely account for the pervasiveness of positive heterosis. Heterosis could almost be viewed as the measurement of the curvature of the GP relationship.

Our results may have practical consequences for heterosis prediction, a major issue in plant and animal breeding. A classical predictor of heterosis is the genetic distance between the parents, but the quality of the prediction depends markedly on the traits, on the genetic history of the material and on the range of genetic distances considered [126,142,155–157]. Our results show that better predictions of heterosis could be achieved by combining parental information at two phenotypic levels, the trait under study and the candidate underlying factors. Omics methods allow thousands of transcripts, proteins and metabolites to be identified and quantified. Thus we now have access to a plethora of putative predictors [158] to be incorporated in non-linear statistical models. The high-dimensionality is challenging, but relevant variables could be selected by using both biological knowledge of the trait under study (e.g. [159]) and recently developed regularization methods [160,161].

## Acknowledgments

We dedicate this work to Pr. André Gallais, who brought us his impressive knowledge of heterosis. We thank Dr. Catherine Damerval, Dr. Michel Zivy, Dr. Mélisande Blein-Nicolas, Dr. Warren Albertin, Dr. Philippe Marullo and Dr. François Vasseur for exciting discussions on heterosis and for useful suggestions for improving the manuscript. We also thank Marianyela Petrizzelli for her help in checking the mathematical developments and Dr. Hélène Citerne for English corrections.

This work was supported by the ANR program “ALIA” HeterosYeast ANR-08-ALIA-9.

### Author Contributions

- **Conceptualization:** Julie B. Fiévet, Christine Dillmann, Dominique de Vienne.
- **Formal Analysis:** Dominique de Vienne.
- **Funding Acquisition:** Christine Dillmann, Dominique de Vienne.
- **Investigation:** Julie B. Fiévet, Thibault Nidelet, Dominique de Vienne.
- **Methodology:** Julie B. Fiévet, Thibault Nidelet, Christine Dillmann, Dominique de Vienne.
- **Project Administration:** Christine Dillmann, Dominique de Vienne.
- **Resources:** Julie B. Fiévet, Thibault Nidelet, Dominique de Vienne.
- **Software:** Thibault Nidelet, Dominique de Vienne.
- **Supervision:** Dominique de Vienne.
- **Validation:** Dominique de Vienne.
- **Visualization:** Dominique de Vienne.
- **Writing – Original Draft Preparation:** Dominique de Vienne.
- **Writing – Review & Editing:** Julie B. Fiévet, Thibault Nidelet, Christine Dillmann, Dominique de Vienne.

## Supporting information

**The geometry of heterosis**, by Julie B. Fiévet, Thibault Nidelet, Christine Dillmann & Dominique de Vienne.

## S1 Appendix **Constrained** *E*_tot_**: consequence on the** *c*_v_**’s of enzyme concentrations.**

Consider the concentration *x*_*j*_ of enzyme *j* before constraint on *E*_tot_. Its expectation is E(*x*_*j*_) = *µ_j_* and its variance:

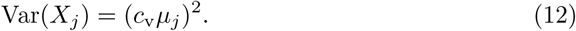

Consider now the variable 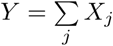. Its expectation is:

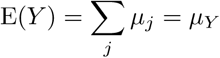

and its variance:

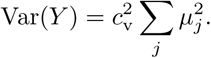

The variable associated with the concentration of enzyme *i* upon constraint on *E*_tot_ is *Z_i_* = *X_i_/Y*. Because *Y* includes *X_i_*, these two variables are not independent. Using the Delta method [162], we can write:

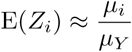

and

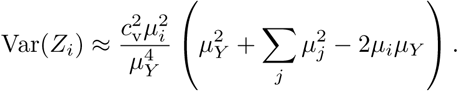

Using Eq 12, we get:

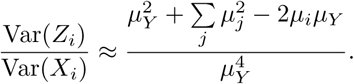

We see that:

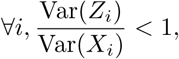

which means that the constraint decreases the variance of every enzyme concentration. Regarding the coefficient of variation *c_i_* of *Z_i_*, we have:

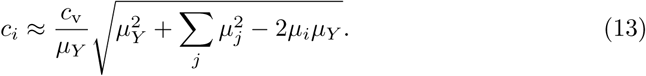

Thus *c_i_/c*_v_ = 1 if 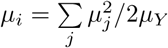, otherwise *c_i_/c*_v_ *<* 1 (resp. *c_i_/c*_v_ *>* 1) when 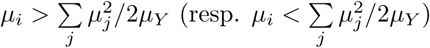. In other words the constraint changes the coefficients of variation in an inverse relation to the enzyme concentrations: the coefficient of variation of the most (resp. less) abundant enzymes will decrease (resp. increase) under constrained *E*_tot_. This theoretical prediction is consistent with observations. For instance, when *c*_v_ = 0.6 in the four-enzyme system, the theoretical coefficients of variation after constraint were 0.63, 0.58, 0.30 and 0.71, for PGI, PFK, FBA and TPI, respectively, and we observed 0.61, 0.54, 0.31 and 0.76. The mean was 0.56, which is close to 0.6.

## S2 Appendix **Comparing initial and actual** *c*_v_**’s of enzyme concentrations in the glycolysis/fermentation system.**

In our simulations of the glycolysis/fermentation network, the steady state was not necessarily reached, and depended on enzyme concentration values. If one member of the triplet parents-hybrid did not reach a steady state, the triplet was excluded, which was all the more frequent when *c*_v_’s were high. The following table shows the percentage of successful simulations under different *c*_v_ values, without and with contraint on *E*_tot_.

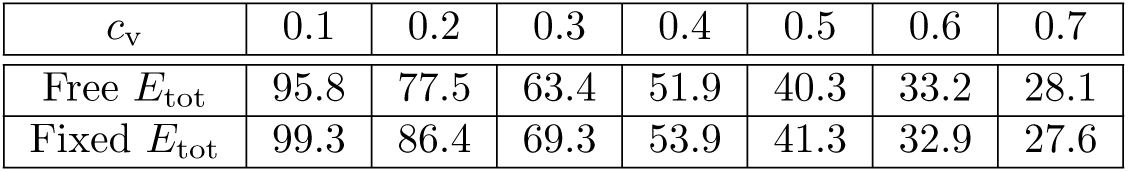

Once 10 000 successful simulations were obtained for each condition, we examined the histograms of glucose flux and observed that there were null values, particularly for the highest *c*_v_’s. The triplets involved were elimitated, so that the total number of simulations finally retained varied according to the *c*_v_ and the presence/absence of constraint:

| $c_v$ | 0.1 | 0.2 | 0.3 | 0.4 | 0.5 | 0.6 | 0.7 |
| --- | --- | --- | --- | --- | --- | --- | --- |
| Free $E_{\text{tot}}$ | 10 000 | 9 998 | 9 727 | 8 634 | 6 738 | 4 812 | 3 162 |
| Fixed $E_{\text{tot}}$ | 10 000 | 10 000 | 10 000 | 9 899 | 9 340 | 8 202 | 6 581 |

Given the high number of excluded simulations for the highest *c*_v_’s, we checked for possible biases in the *a posteriori* distributions of enzyme concentrations. This was actually found to be the case, as shown S9 Fig. With free *E*_tot_, the mean concentrations of PFK, FBA and ADH, and to a lesser extent of PGK, were higher than the reference concentrations, while the concentration of HK was lower, this effect being positively related to the initial *c*_v_ values (S9 FigA). This means that, frequently, the lowest concentrations of some enzymes (the highest for HK) did not allow the steady state to be reached or resulted in null fluxes, and so the corresponding triplets were excluded. As a consequence, means and variances were modified and *c*_v_’s changed accordingly, the *c*_v_ of HK being the most reduced for high initial *c*_v_ values (S9 FigB).

With fixed *E*_tot_, the *a posteriori* means were markedly different from the reference concentrations for the highest *c*_v_’s (S9 Fig C). This is likely due to HK, which is by far the most abundant enzyme (77.5 % of the total concentration): whenever a high HK value was drawn, application of the constraint reduced the concentration of the other enzymes, which can prevent the steady state from being reached or can lead to null fluxes. Therefore the mean concentration of HK was reduced and the concentrations of the other enzymes were increased, the variances of the concentrations being decreased in both cases. The resulting *c*_v_’s are shown S9 FigE, in comparison with the theoretical values that are expected in the case of constraint (Eq 13 and S9 FigD): again, discrepancies were observed for the highest *c*_v_’s and depended on the enzyme. In these cases, observed *c*_v_’s were generally lower than theoretical *c*_v_’s.

---

**S1 Table.** Enzyme concentrations (mg.L^−1^) of the parents of the 61 *in vitro* crosses. PGI, phosphoglucose isomerase; PFK, phosphofructokinase; FBA, fructose-1,6-bisphosphate aldolase; TPI, triosephosphate isomerase.

| Cross | Parent 1 |  |  |  |  | Parent 2 |  |  |  |  |
| --- | --- | --- | --- | --- | --- | --- | --- | --- | --- | --- |
| # | # | PGI | PFK | FBA | TPI | # | PGI | PFK | FBA | TPI |
| 1 | 24 | 70 | 5 | 5 | 21.9 | 40 | 25 | 70 | 2 | 4.9 |
| 2 | 30 | 40 | 35 | 5 | 21.9 | 38 | 35 | 60 | 5 | 1.9 |
| 3 | 27 | 55 | 15 | 12 | 19.9 | 38 | 35 | 60 | 5 | 1.9 |
| 4 | 25 | 20 | 10 | 5 | 66.9 | 36 | 25 | 50 | 12 | 14.9 |
| 5 | 27 | 55 | 15 | 12 | 19.9 | 35 | 15 | 50 | 5 | 31.9 |
| 6 | 29 | 10 | 20 | 15 | 56.9 | 30 | 40 | 35 | 5 | 21.9 |
| 7 | 24 | 70 | 5 | 5 | 21.9 | 26 | 40 | 10 | 40 | 11.9 |
| 8 | 14 | 4.23 | 2.62 | 76.42 | 18.62 | 16 | 3.72 | 1.95 | 86.61 | 9.61 |
| 9 | 16 | 3.72 | 1.95 | 86.61 | 9.61 | 13 | 5.79 | 3.3 | 76.42 | 16.38 |
| 10 | 15 | 9.4 | 2.58 | 86.61 | 3.31 | 23 | 3.4 | 2.81 | 86.61 | 9.08 |
| 11 | 14 | 4.23 | 2.62 | 76.42 | 18.62 | 23 | 3.4 | 2.81 | 86.61 | 9.08 |
| 12 | 23 | 3.4 | 2.81 | 86.61 | 9.08 | 17 | 7.36 | 3.21 | 86.61 | 4.72 |
| 13 | 17 | 7.36 | 3.21 | 86.61 | 4.72 | 22 | 9.88 | 3.73 | 86.61 | 1.67 |
| 14 | 40 | 25 | 70 | 2 | 4.9 | 28 | 15 | 15 | 55 | 16.9 |
| 15 | 1 | 39.32 | 12.38 | 25.47 | 24.71 | 2 | 51.97 | 14.22 | 25.47 | 10.23 |
| 16 | 15 | 9.4 | 2.58 | 86.61 | 3.31 | 8 | 3.33 | 5.75 | 66.23 | 26.58 |
| 17 | 23 | 3.4 | 2.81 | 86.61 | 9.08 | 12 | 5.92 | 5.25 | 76.42 | 14.3 |
| 18 | 13 | 5.79 | 3.3 | 76.42 | 16.38 | 8 | 3.33 | 5.75 | 66.23 | 26.58 |
| 19 | 32 | 15 | 40 | 45 | 1.9 | 37 | 25 | 50 | 20 | 6.9 |
| 20 | 13 | 5.79 | 3.3 | 76.42 | 16.38 | 19 | 29.71 | 6.62 | 56.04 | 9.53 |
| 21 | 17 | 7.36 | 3.21 | 86.61 | 4.72 | 8 | 3.33 | 5.75 | 66.23 | 26.58 |
| 22 | 14 | 4.23 | 2.62 | 76.42 | 18.62 | 20 | 13.29 | 8.21 | 56.04 | 24.36 |
| 23 | 17 | 7.36 | 3.21 | 86.61 | 4.72 | 19 | 29.71 | 6.62 | 56.04 | 9.53 |
| 24 | 2 | 51.97 | 14.22 | 25.47 | 10.23 | 5 | 15.66 | 23.52 | 45.85 | 16.86 |
| 25 | 15 | 9.4 | 2.58 | 86.61 | 3.31 | 18 | 37.9 | 12.6 | 35.66 | 15.73 |
| 26 | 15 | 9.4 | 2.58 | 86.61 | 3.31 | 1 | 39.32 | 12.38 | 25.47 | 24.71 |
| 27 | 22 | 9.88 | 3.73 | 86.61 | 1.67 | 21 | 20.6 | 7.18 | 66.23 | 7.89 |
| 28 | 14 | 4.23 | 2.62 | 76.42 | 18.62 | 2 | 51.97 | 14.22 | 25.47 | 10.23 |
| 29 | 18 | 37.9 | 12.6 | 35.66 | 15.73 | 7 | 6.79 | 20.43 | 56.04 | 18.63 |
| 30 | 13 | 5.79 | 3.3 | 76.42 | 16.38 | 1 | 39.32 | 12.38 | 25.47 | 24.71 |
| 31 | 2 | 51.97 | 14.22 | 25.47 | 10.23 | 8 | 3.33 | 5.75 | 66.23 | 26.58 |
| 32 | 19 | 29.71 | 6.62 | 56.04 | 9.53 | 18 | 37.9 | 12.6 | 35.66 | 15.73 |
| 33 | 12 | 5.92 | 5.25 | 76.42 | 14.3 | 1 | 39.32 | 12.38 | 25.47 | 24.71 |
| 34 | 19 | 29.71 | 6.62 | 56.04 | 9.53 | 20 | 13.29 | 8.21 | 56.04 | 24.36 |
| 35 | 10 | 4.24 | 25.75 | 66.23 | 5.67 | 4 | 32.45 | 28.74 | 35.66 | 5.03 |
| 36 | 11 | 8.44 | 9.53 | 66.23 | 17.69 | 4 | 32.45 | 28.74 | 35.66 | 5.03 |
| 37 | 13 | 5.79 | 3.3 | 76.42 | 16.38 | 4 | 32.45 | 28.74 | 35.66 | 5.03 |
| 38 | 20 | 13.29 | 8.21 | 56.04 | 24.36 | 8 | 3.33 | 5.75 | 66.23 | 26.58 |
| 39 | 11 | 8.44 | 9.53 | 66.23 | 17.69 | 6 | 15.55 | 30.53 | 45.85 | 9.96 |
| 40 | 20 | 13.29 | 8.21 | 56.04 | 24.36 | 9 | 9.01 | 8.8 | 66.23 | 17.85 |
| 41 | 5 | 15.66 | 23.52 | 45.85 | 16.86 | 4 | 32.45 | 28.74 | 35.66 | 5.03 |
| 42 | 10 | 4.24 | 25.75 | 66.23 | 5.67 | 6 | 15.55 | 30.53 | 45.85 | 9.96 |

| *Continued from the previous page* |  |  |  |  |  |  |  |  |  |  |
| --- | --- | --- | --- | --- | --- | --- | --- | --- | --- | --- |
| **Cross** | **Parent 1** |  |  |  |  | **Parent 2** |  |  |  |  |
| **#** | **#** | **PGI** | **PFK** | **FBA** | **TPI** | **#** | **PGI** | **PFK** | **FBA** | **TPI** |
| 43 | 7 | 6.79 | 20.43 | 56.04 | 18.63 | 6 | 15.55 | 30.53 | 45.85 | 9.96 |
| 44 | 16 | 3.72 | 1.95 | 86.61 | 9.61 | 7 | 6.79 | 20.43 | 56.04 | 18.63 |
| 45 | 12 | 5.92 | 5.25 | 76.42 | 14.3 | 4 | 32.45 | 28.74 | 35.66 | 5.03 |
| 46 | 18 | 37.9 | 12.6 | 35.66 | 15.73 | 9 | 9.01 | 8.8 | 66.23 | 17.85 |
| 47 | 5 | 15.66 | 23.52 | 45.85 | 16.86 | 7 | 6.79 | 20.43 | 56.04 | 18.63 |
| 48 | 15 | 9.4 | 2.58 | 86.61 | 3.31 | 10 | 4.24 | 25.75 | 66.23 | 5.67 |
| 49 | 12 | 5.92 | 5.25 | 76.42 | 14.3 | 3 | 12.2 | 27.64 | 35.66 | 26.39 |
| 50 | 11 | 8.44 | 9.53 | 66.23 | 17.69 | 2 | 51.97 | 14.22 | 25.47 | 10.23 |
| 51 | 16 | 3.72 | 1.95 | 86.61 | 9.61 | 6 | 15.55 | 30.53 | 45.85 | 9.96 |
| 52 | 3 | 12.2 | 27.64 | 35.66 | 26.39 | 9 | 9.01 | 8.8 | 66.23 | 17.85 |
| 53 | 5 | 15.66 | 23.52 | 45.85 | 16.86 | 9 | 9.01 | 8.8 | 66.23 | 17.85 |
| 54 | 17 | 7.36 | 3.21 | 86.61 | 4.72 | 1 | 39.32 | 12.38 | 25.47 | 24.71 |
| 55 | 21 | 20.6 | 7.18 | 66.23 | 7.89 | 6 | 15.55 | 30.53 | 45.85 | 9.96 |
| 56 | 21 | 20.6 | 7.18 | 66.23 | 7.89 | 3 | 12.2 | 27.64 | 35.66 | 26.39 |
| 57 | 21 | 20.6 | 7.18 | 66.23 | 7.89 | 5 | 15.66 | 23.52 | 45.85 | 16.86 |
| 58 | 10 | 4.24 | 25.75 | 66.23 | 5.67 | 22 | 9.88 | 3.73 | 86.61 | 1.67 |
| 59 | 10 | 4.24 | 25.75 | 66.23 | 5.67 | 12 | 5.92 | 5.25 | 76.42 | 14.3 |
| 60 | 22 | 9.88 | 3.73 | 86.61 | 1.67 | 3 | 12.2 | 27.64 | 35.66 | 26.39 |
| 61 | 17 | 7.36 | 3.21 | 86.61 | 4.72 | 5 | 15.66 | 23.52 | 45.85 | 16.86 |

**S2 Table.**
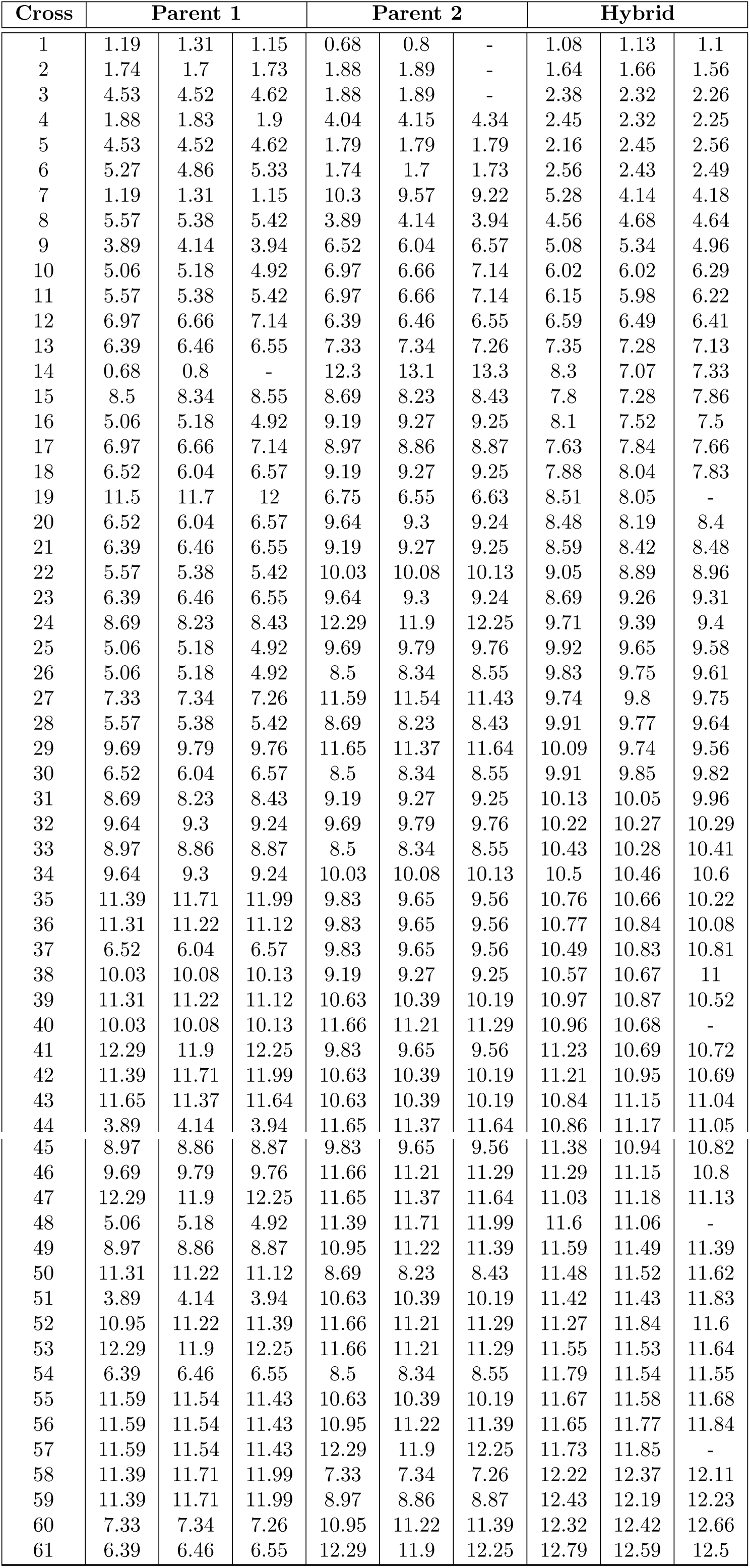
Flux values (*µ*M.s^−1^) in parent and hybrid replicates.

| Cross | Parent 1 |  |  | Parent 2 |  |  | Hybrid |  |  |
| --- | --- | --- | --- | --- | --- | --- | --- | --- | --- |
| 1 | 1.19 | 1.31 | 1.15 | 0.68 | 0.8 | - | 1.08 | 1.13 | 1.1 |
| 2 | 1.74 | 1.7 | 1.73 | 1.88 | 1.89 | - | 1.64 | 1.66 | 1.56 |
| 3 | 4.53 | 4.52 | 4.62 | 1.88 | 1.89 | - | 2.38 | 2.32 | 2.26 |
| 4 | 1.88 | 1.83 | 1.9 | 4.04 | 4.15 | 4.34 | 2.45 | 2.32 | 2.25 |
| 5 | 4.53 | 4.52 | 4.62 | 1.79 | 1.79 | 1.79 | 2.16 | 2.45 | 2.56 |
| 6 | 5.27 | 4.86 | 5.33 | 1.74 | 1.7 | 1.73 | 2.56 | 2.43 | 2.49 |
| 7 | 1.19 | 1.31 | 1.15 | 10.3 | 9.57 | 9.22 | 5.28 | 4.14 | 4.18 |
| 8 | 5.57 | 5.38 | 5.42 | 3.89 | 4.14 | 3.94 | 4.56 | 4.68 | 4.64 |
| 9 | 3.89 | 4.14 | 3.94 | 6.52 | 6.04 | 6.57 | 5.08 | 5.34 | 4.96 |
| 10 | 5.06 | 5.18 | 4.92 | 6.97 | 6.66 | 7.14 | 6.02 | 6.02 | 6.29 |
| 11 | 5.57 | 5.38 | 5.42 | 6.97 | 6.66 | 7.14 | 6.15 | 5.98 | 6.22 |
| 12 | 6.97 | 6.66 | 7.14 | 6.39 | 6.46 | 6.55 | 6.59 | 6.49 | 6.41 |
| 13 | 6.39 | 6.46 | 6.55 | 7.33 | 7.34 | 7.26 | 7.35 | 7.28 | 7.13 |
| 14 | 0.68 | 0.8 | - | 12.3 | 13.1 | 13.3 | 8.3 | 7.07 | 7.33 |
| 15 | 8.5 | 8.34 | 8.55 | 8.69 | 8.23 | 8.43 | 7.8 | 7.28 | 7.86 |
| 16 | 5.06 | 5.18 | 4.92 | 9.19 | 9.27 | 9.25 | 8.1 | 7.52 | 7.5 |
| 17 | 6.97 | 6.66 | 7.14 | 8.97 | 8.86 | 8.87 | 7.63 | 7.84 | 7.66 |
| 18 | 6.52 | 6.04 | 6.57 | 9.19 | 9.27 | 9.25 | 7.88 | 8.04 | 7.83 |
| 19 | 11.5 | 11.7 | 12 | 6.75 | 6.55 | 6.63 | 8.51 | 8.05 | - |
| 20 | 6.52 | 6.04 | 6.57 | 9.64 | 9.3 | 9.24 | 8.48 | 8.19 | 8.4 |
| 21 | 6.39 | 6.46 | 6.55 | 9.19 | 9.27 | 9.25 | 8.59 | 8.42 | 8.48 |
| 22 | 5.57 | 5.38 | 5.42 | 10.03 | 10.08 | 10.13 | 9.05 | 8.89 | 8.96 |
| 23 | 6.39 | 6.46 | 6.55 | 9.64 | 9.3 | 9.24 | 8.69 | 9.26 | 9.31 |
| 24 | 8.69 | 8.23 | 8.43 | 12.29 | 11.9 | 12.25 | 9.71 | 9.39 | 9.4 |
| 25 | 5.06 | 5.18 | 4.92 | 9.69 | 9.79 | 9.76 | 9.92 | 9.65 | 9.58 |
| 26 | 5.06 | 5.18 | 4.92 | 8.5 | 8.34 | 8.55 | 9.83 | 9.75 | 9.61 |
| 27 | 7.33 | 7.34 | 7.26 | 11.59 | 11.54 | 11.43 | 9.74 | 9.8 | 9.75 |
| 28 | 5.57 | 5.38 | 5.42 | 8.69 | 8.23 | 8.43 | 9.91 | 9.77 | 9.64 |
| 29 | 9.69 | 9.79 | 9.76 | 11.65 | 11.37 | 11.64 | 10.09 | 9.74 | 9.56 |
| 30 | 6.52 | 6.04 | 6.57 | 8.5 | 8.34 | 8.55 | 9.91 | 9.85 | 9.82 |
| 31 | 8.69 | 8.23 | 8.43 | 9.19 | 9.27 | 9.25 | 10.13 | 10.05 | 9.96 |
| 32 | 9.64 | 9.3 | 9.24 | 9.69 | 9.79 | 9.76 | 10.22 | 10.27 | 10.29 |
| 33 | 8.97 | 8.86 | 8.87 | 8.5 | 8.34 | 8.55 | 10.43 | 10.28 | 10.41 |
| 34 | 9.64 | 9.3 | 9.24 | 10.03 | 10.08 | 10.13 | 10.5 | 10.46 | 10.6 |
| 35 | 11.39 | 11.71 | 11.99 | 9.83 | 9.65 | 9.56 | 10.76 | 10.66 | 10.22 |
| 36 | 11.31 | 11.22 | 11.12 | 9.83 | 9.65 | 9.56 | 10.77 | 10.84 | 10.08 |
| 37 | 6.52 | 6.04 | 6.57 | 9.83 | 9.65 | 9.56 | 10.49 | 10.83 | 10.81 |
| 38 | 10.03 | 10.08 | 10.13 | 9.19 | 9.27 | 9.25 | 10.57 | 10.67 | 11 |
| 39 | 11.31 | 11.22 | 11.12 | 10.63 | 10.39 | 10.19 | 10.97 | 10.87 | 10.52 |
| 40 | 10.03 | 10.08 | 10.13 | 11.66 | 11.21 | 11.29 | 10.96 | 10.68 | - |
| 41 | 12.29 | 11.9 | 12.25 | 9.83 | 9.65 | 9.56 | 11.23 | 10.69 | 10.72 |
| 42 | 11.39 | 11.71 | 11.99 | 10.63 | 10.39 | 10.19 | 11.21 | 10.95 | 10.69 |
| 43 | 11.65 | 11.37 | 11.64 | 10.63 | 10.39 | 10.19 | 10.84 | 11.15 | 11.04 |
| 44 | 3.89 | 4.14 | 3.94 | 11.65 | 11.37 | 11.64 | 10.86 | 11.17 | 11.05 |

| *Continued from the previous page* |  |  |  |  |  |  |  |  |  |
| --- | --- | --- | --- | --- | --- | --- | --- | --- | --- |
| **Cross** | **Parent 1** |  |  | **Parent 2** |  |  | **Hybrid** |  |  |
| 45 | 8.97 | 8.86 | 8.87 | 9.83 | 9.65 | 9.56 | 11.38 | 10.94 | 10.82 |
| 46 | 9.69 | 9.79 | 9.76 | 11.66 | 11.21 | 11.29 | 11.29 | 11.15 | 10.8 |
| 47 | 12.29 | 11.9 | 12.25 | 11.65 | 11.37 | 11.64 | 11.03 | 11.18 | 11.13 |
| 48 | 5.06 | 5.18 | 4.92 | 11.39 | 11.71 | 11.99 | 11.6 | 11.06 | - |
| 49 | 8.97 | 8.86 | 8.87 | 10.95 | 11.22 | 11.39 | 11.59 | 11.49 | 11.39 |
| 50 | 11.31 | 11.22 | 11.12 | 8.69 | 8.23 | 8.43 | 11.48 | 11.52 | 11.62 |
| 51 | 3.89 | 4.14 | 3.94 | 10.63 | 10.39 | 10.19 | 11.42 | 11.43 | 11.83 |
| 52 | 10.95 | 11.22 | 11.39 | 11.66 | 11.21 | 11.29 | 11.27 | 11.84 | 11.6 |
| 53 | 12.29 | 11.9 | 12.25 | 11.66 | 11.21 | 11.29 | 11.55 | 11.53 | 11.64 |
| 54 | 6.39 | 6.46 | 6.55 | 8.5 | 8.34 | 8.55 | 11.79 | 11.54 | 11.55 |
| 55 | 11.59 | 11.54 | 11.43 | 10.63 | 10.39 | 10.19 | 11.67 | 11.58 | 11.68 |
| 56 | 11.59 | 11.54 | 11.43 | 10.95 | 11.22 | 11.39 | 11.65 | 11.77 | 11.84 |
| 57 | 11.59 | 11.54 | 11.43 | 12.29 | 11.9 | 12.25 | 11.73 | 11.85 | - |
| 58 | 11.39 | 11.71 | 11.99 | 7.33 | 7.34 | 7.26 | 12.22 | 12.37 | 12.11 |
| 59 | 11.39 | 11.71 | 11.99 | 8.97 | 8.86 | 8.87 | 12.43 | 12.19 | 12.23 |
| 60 | 7.33 | 7.34 | 7.26 | 10.95 | 11.22 | 11.39 | 12.32 | 12.42 | 12.66 |
| 61 | 6.39 | 6.46 | 6.55 | 12.29 | 11.9 | 12.25 | 12.79 | 12.59 | 12.5 |

**S3 Table.** Parameters of the variable enzymes. *V*_max_: maximal velocity, *k*_cat_: catalytic constant, Mr: molecular mass, *E*_ref_: reference concentration. For the simulations, we used *E*_ref_*/*5 (see Materials and methods).

| Enzyme | $V_{\max}$<br>mMol.min<sup>-1</sup> | $k_{\text{cat}}$<br>min<sup>-1</sup> | Mr<br>g/mMol | $E_{\text{ref}}$<br>mg |
| --- | --- | --- | --- | --- |
| HK (E.C. 2.7.1.1) | 236.70 | 105.6 | 55 | 123281.25 |
| PGI (E.C. 5.3.1.9) | 1056.00 | 84600.0 | 120 | 1466.67 |
| PFK (E.C. 2.7.1.11) | 110.00 | 554400.0 | 790 | 156.75 |
| PGK (E.C. 2.7.2.3) | 1288.00 | 21240.0 | 50 | 3032.02 |
| FBA (E.C. 4.1.2.13) | 94.69 | 12500.0 | 80 | 606.02 |
| PGM (E.C. 5.4.2.11) | 2585.00 | 29400.0 | 112.4 | 9882.79 |
| ENO (E.C. 4.2.1.11) | 201.60 | 4680.0 | 90 | 3876.92 |
| PYK (E.C. 2.7.1.40) | 1000.00 | 13920.0 | 216 | 15517.24 |
| PDC (E.C. 4.1.1.1) | 857.80 | 232000.0 | 250 | 924.35 |
| ADH (E.C. 1.1.1.1) | 209.50 | 47166.0 | 60 | 266.51 |
| G3PDH (E.C. 1.1.1.8) | 47.11 | 20700.0 | 70 | 159.31 |

**S4 Table.**
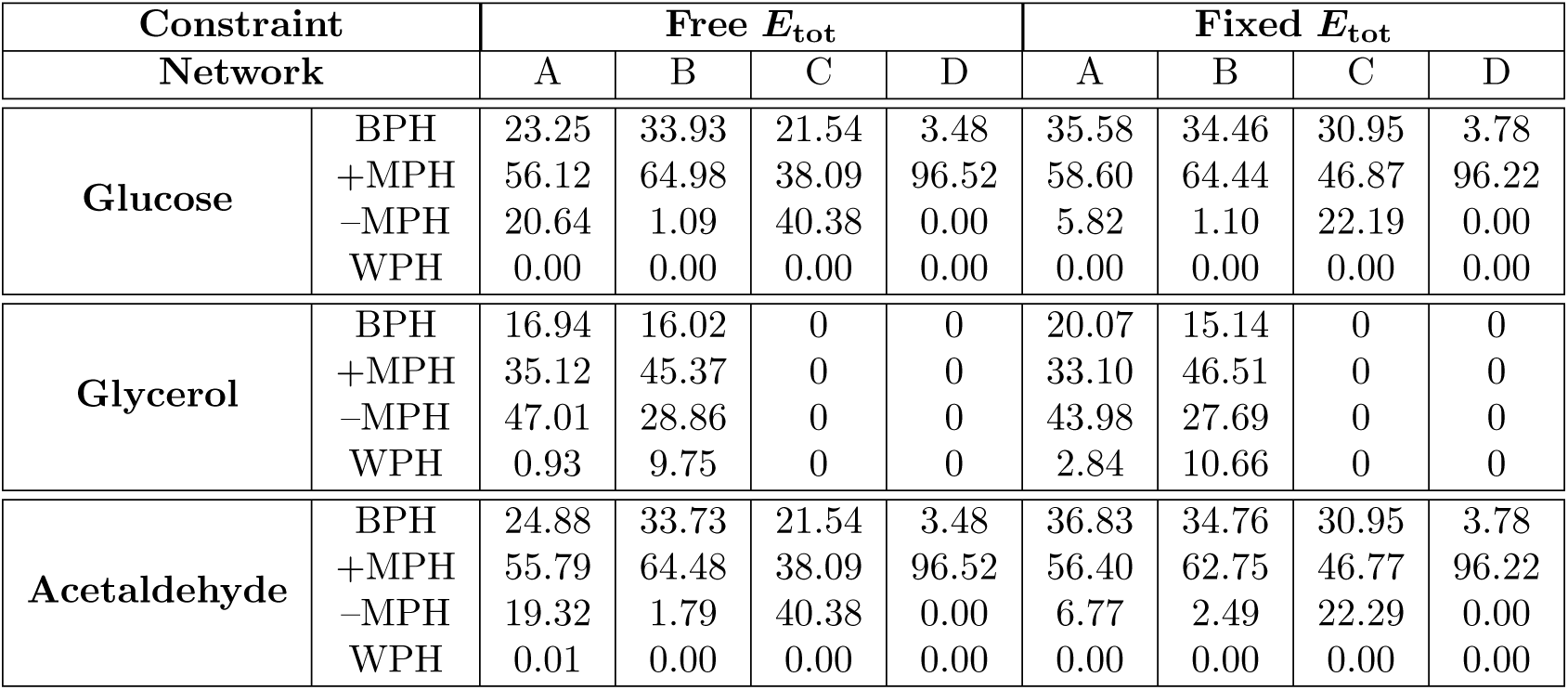
Percentages of the four types of inheritance for the three fluxes in different networks. A: Normal network. B: With constant hexokinase. C: No glycerol branch. D: With constant hexokinase and no glycerol branch (see S4 Fig).

| Constraint | | Free $E_{\text{tot}}$ | | | | Fixed $E_{\text{tot}}$ | | | |
| --- | --- | --- | --- | --- | --- | --- | --- | --- | --- |
| Network |  | A | B | C | D | A | B | C | D |
| Glucose | BPH | 23.25 | 33.93 | 21.54 | 3.48 | 35.58 | 34.46 | 30.95 | 3.78 |
|  | +MPH | 56.12 | 64.98 | 38.09 | 96.52 | 58.60 | 64.44 | 46.87 | 96.22 |
|  | -MPH | 20.64 | 1.09 | 40.38 | 0.00 | 5.82 | 1.10 | 22.19 | 0.00 |
|  | WPH | 0.00 | 0.00 | 0.00 | 0.00 | 0.00 | 0.00 | 0.00 | 0.00 |
| Glycerol | BPH | 16.94 | 16.02 | 0 | 0 | 20.07 | 15.14 | 0 | 0 |
|  | +MPH | 35.12 | 45.37 | 0 | 0 | 33.10 | 46.51 | 0 | 0 |
|  | -MPH | 47.01 | 28.86 | 0 | 0 | 43.98 | 27.69 | 0 | 0 |
|  | WPH | 0.93 | 9.75 | 0 | 0 | 2.84 | 10.66 | 0 | 0 |
| Acetaldehyde | BPH | 24.88 | 33.73 | 21.54 | 3.48 | 36.83 | 34.76 | 30.95 | 3.78 |
|  | +MPH | 55.79 | 64.48 | 38.09 | 96.52 | 56.40 | 62.75 | 46.77 | 96.22 |
|  | -MPH | 19.32 | 1.79 | 40.38 | 0.00 | 6.77 | 2.49 | 22.29 | 0.00 |
|  | WPH | 0.01 | 0.00 | 0.00 | 0.00 | 0.00 | 0.00 | 0.00 | 0.00 |

**S1 Fig.**
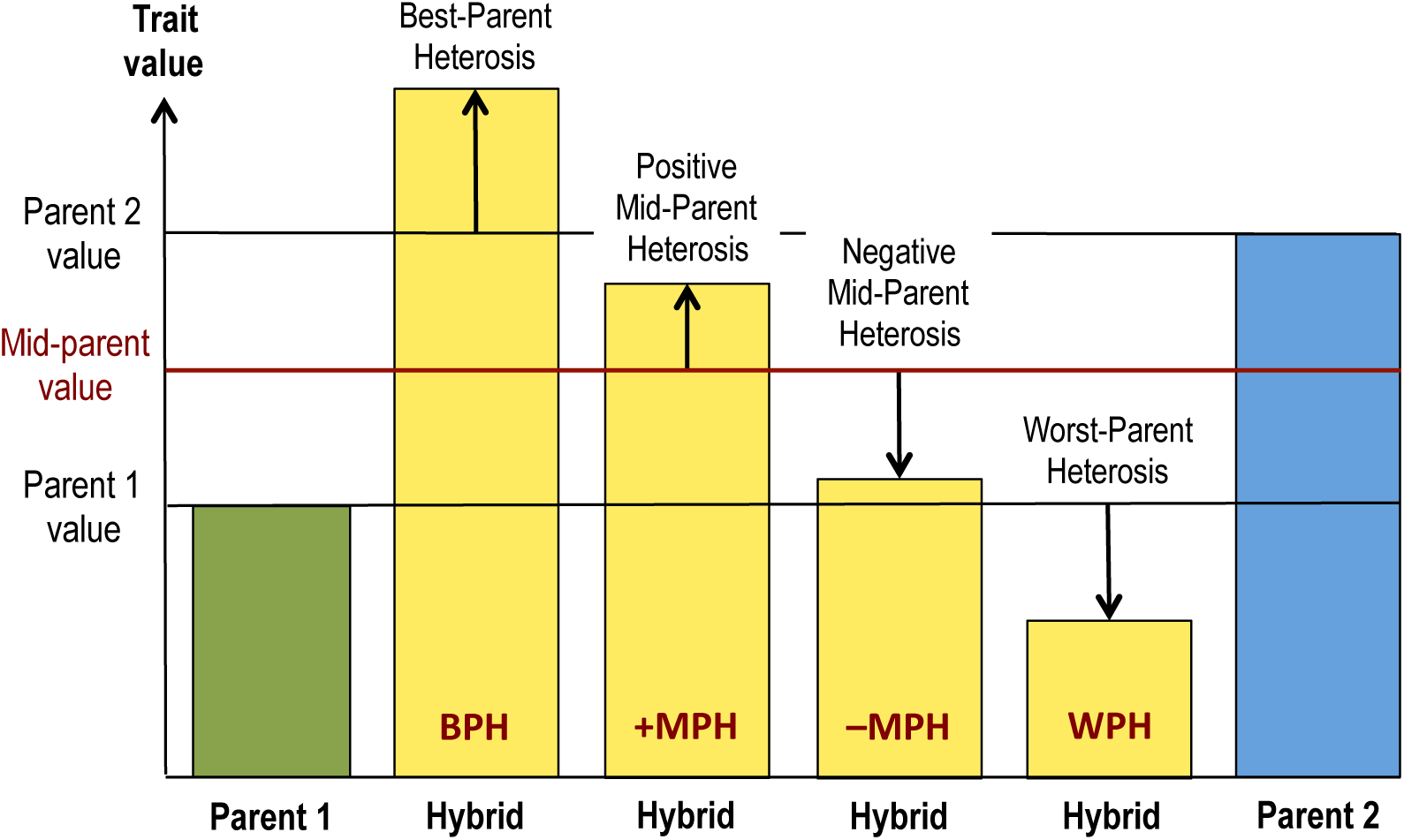
Heterosis types.

**S2 Fig.**
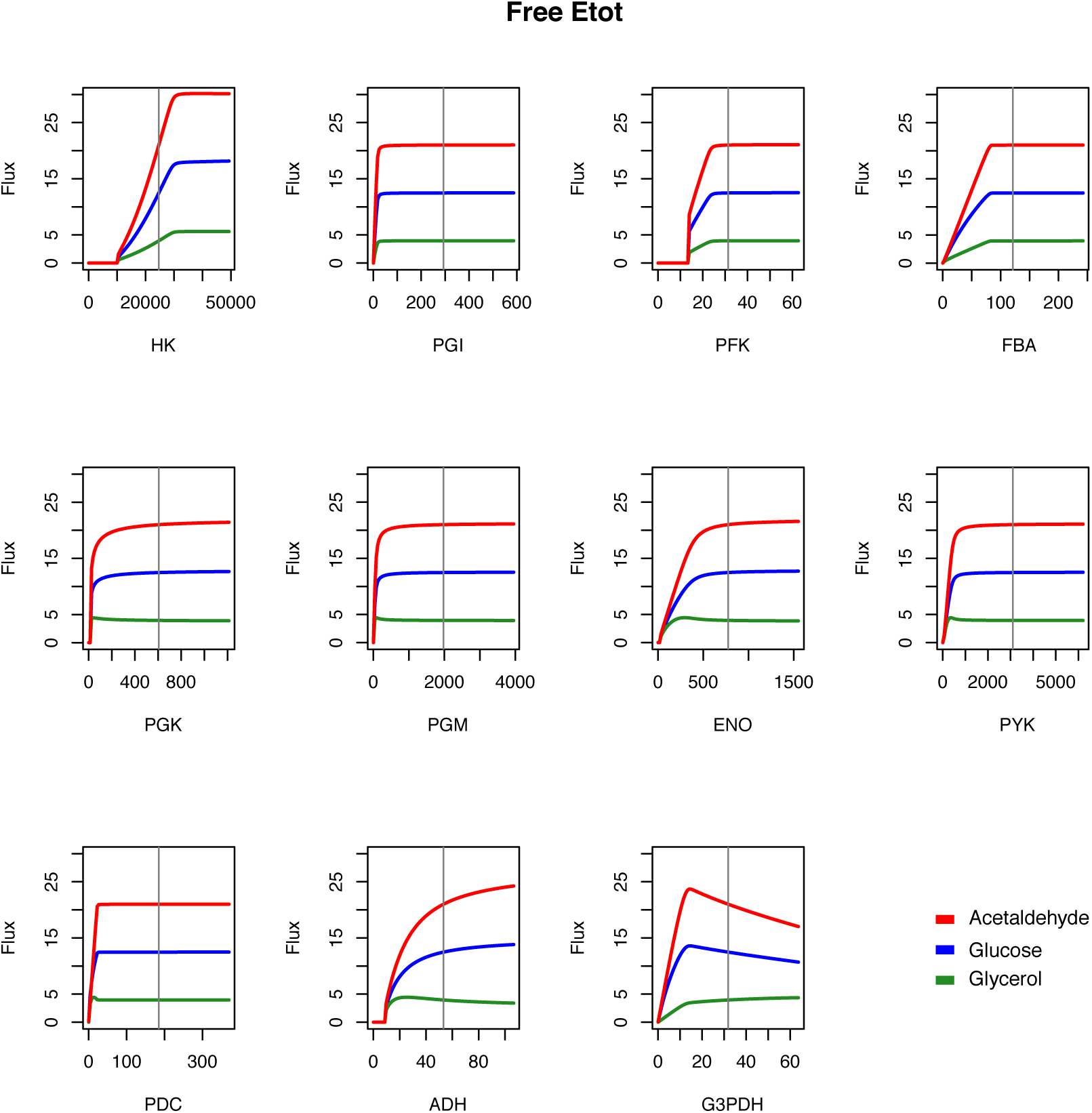
Flux responses to variations of enzyme concentration when *E*_tot_ is free. Enzyme concentrations varied from 0 to twice the reference values chosen for the simulations (see S3 Table), with no constraint on *E*_tot_. Vertical lines indicate reference concentrations.

**S3 Fig.**
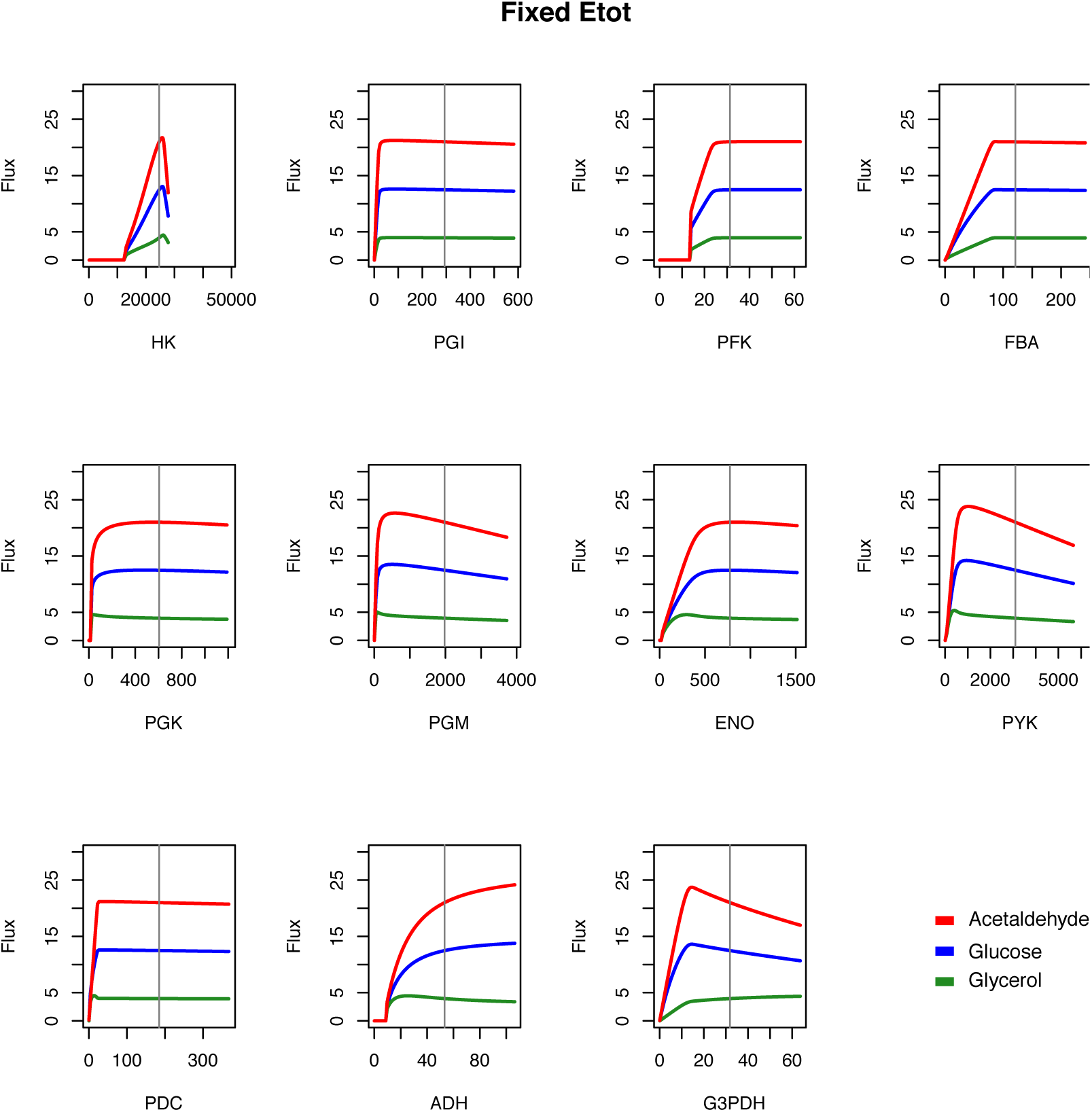
Flux responses to variations of enzyme concentration when *E*_tot_ is fixed. Same as S2 Fig, except that *E*_tot_ was fixed.

**S4 Fig.**
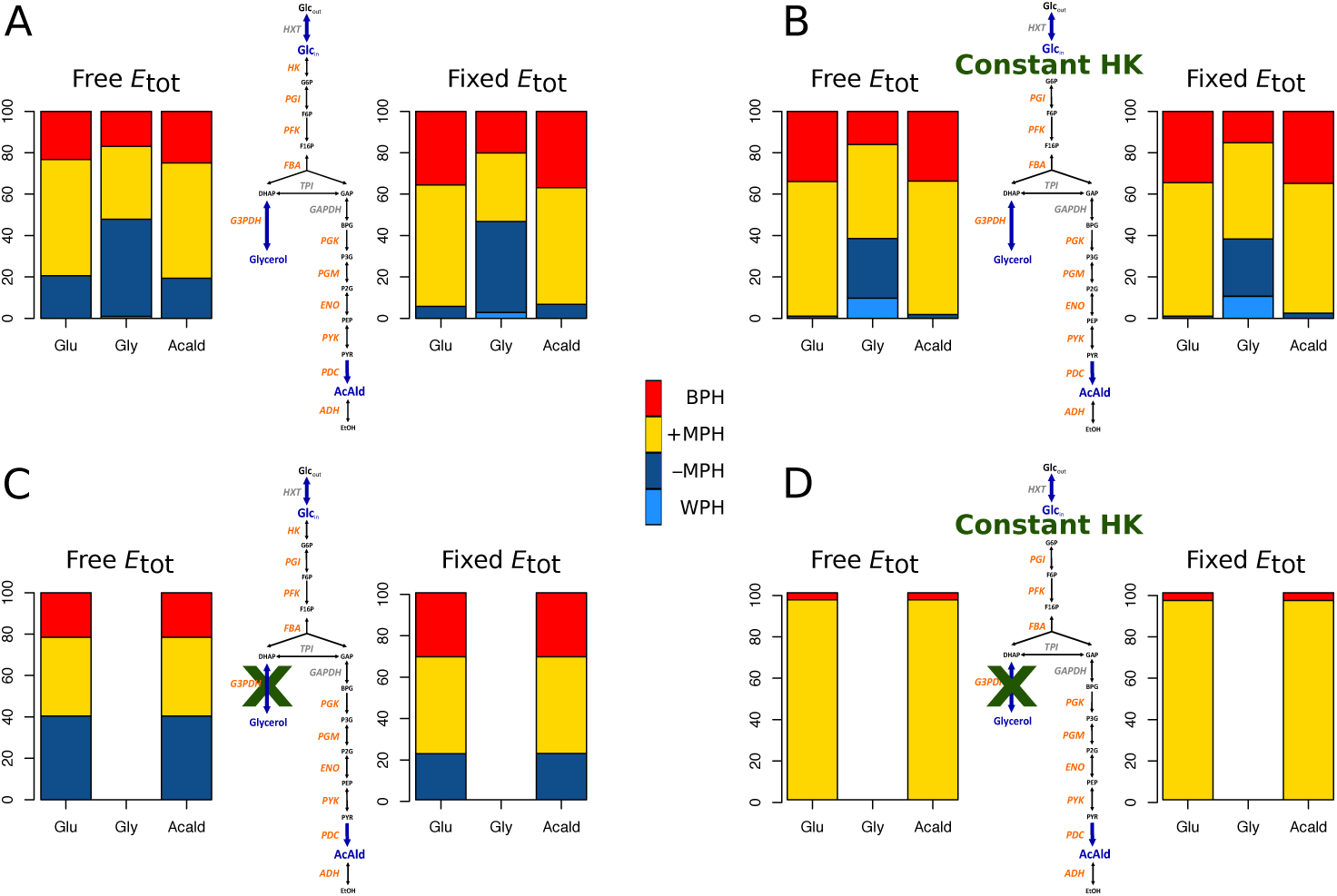
Comparison of the percentages of different types of inheritance with and without convexity in the enzyme-flux relationship. A: Normal network. B: Network with constant hexokinase. C: Network without the glycerol branch. D: Network with constant hexokinase and without the glycerol branch (*c*_v_ = 0.4).

**S5 Fig.**
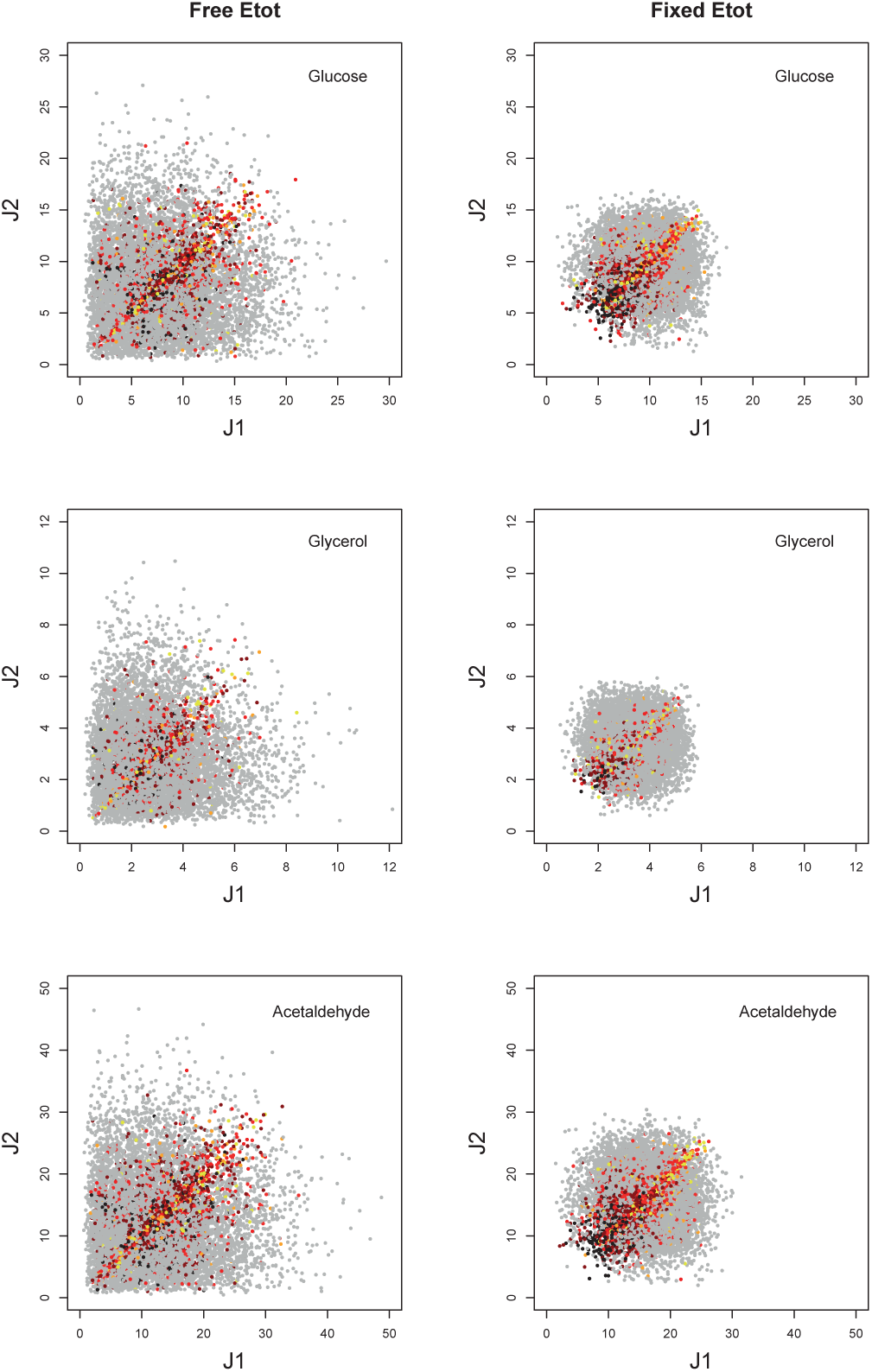
Relationship between parental (*J*_1_ and *J*_2_) and hybrid fluxes. Symbols as in Fig 8. *c*_v_ = 0.4.

**S6 Fig.**
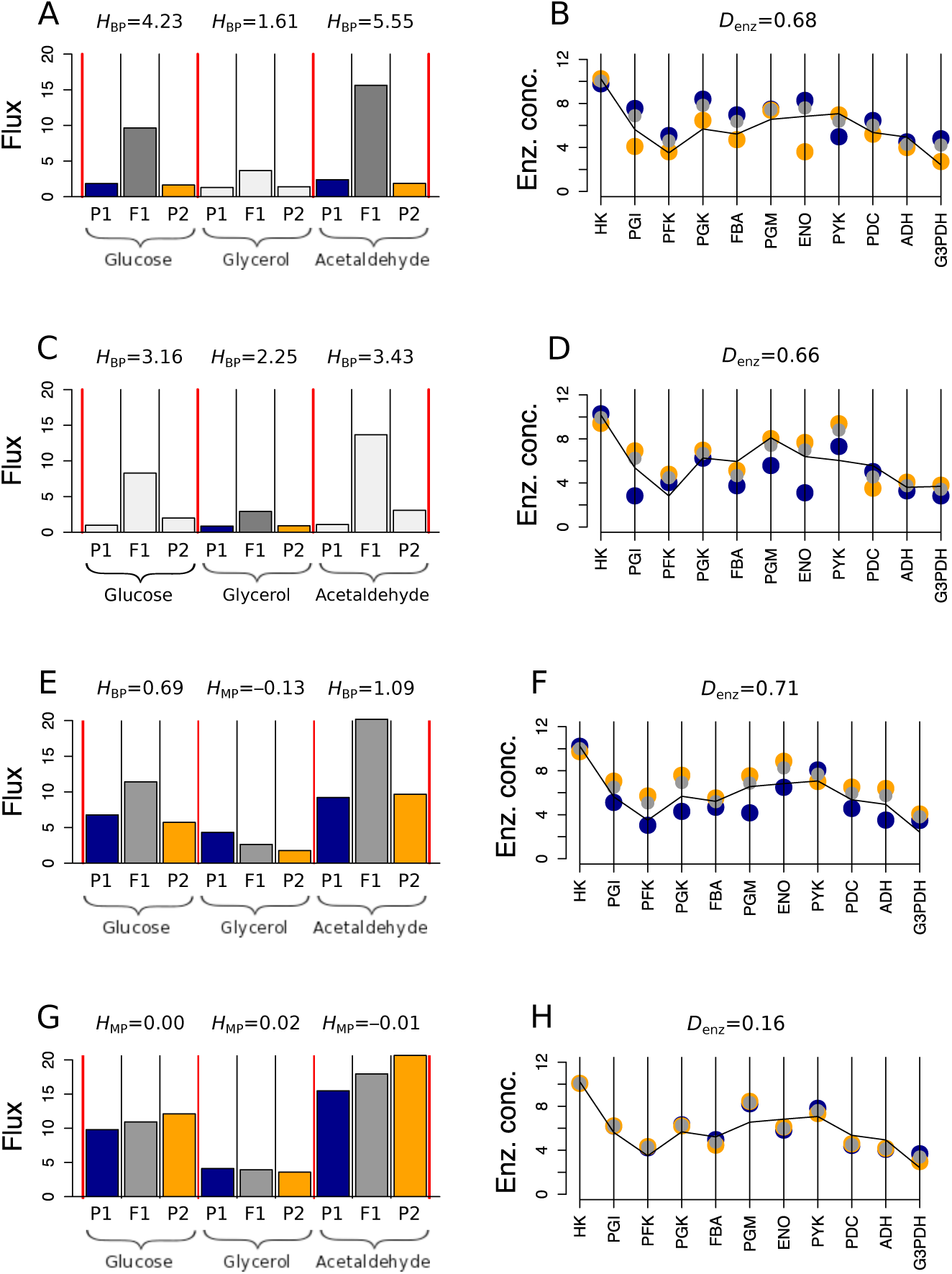
Examples of relationship between inheritance and enzyme concentrations. (*c*_v_ = 0.7, fixed *E*_tot_). A: Parental (blue and orange) and hybrid (gray) fluxes for the cross that displayed the highest heterosis for glucose. This cross also displayed the highest heterosis for acetaldehyde; B: Corresponding enzyme concentrations. C: Fluxes for the cross that displayed the highest heterosis for glycerol. D: Corresponding enzyme concentrations. E: Flux values in the cross between the most distant parents (*D*_enz_ = 0.71). F: Corresponding enzyme concentrations. G: Flux values in the cross between the closest parents (*D*_enz_ = 0.16). H: Corresponding enzyme concentrations. The broken line in B, D, F and H shows the enzyme concentrations of the parent displaying the highest glucose flux value, as a proxy for the optimal concentration distribution. Enzyme concentrations are log-transformed. When there is no BPH (*H*_BP_ *<* 0) we display the index *H*_MP_, the sign of which indicates whether there is positive or negative MPH.

**S7 Fig.**
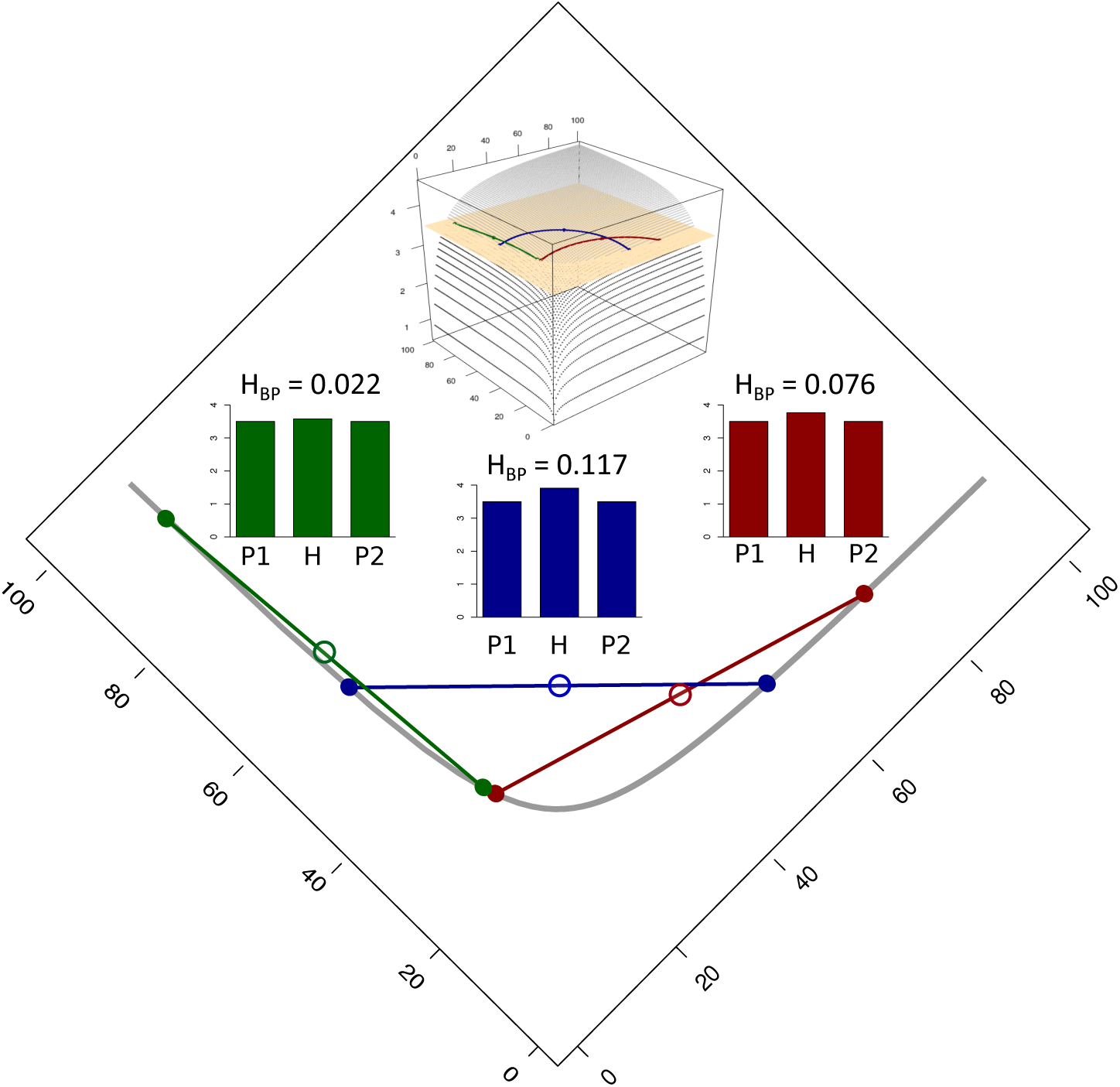
Relationship between heterosis and the position of the parents in the concentration space of two enzymes. Three pairs of parents (green, blue and red) are on the same level curve (points on the gray curve) defined by the horizontal cutting plane shown on the 3D vignette. *D*_enz_ is the same for the three pairs of parents. There is additivity of enzyme concentrations in the hybrids (open circles). BPH is higher when parents are complementary for “high” and “low” concentrations (compare blue with red and green cases). The figure is drawn from the equation: 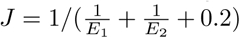 in arbitrary units. The level curve is at *J* = 3.5, and *D_enz_* = 60.

**S8 Fig.**
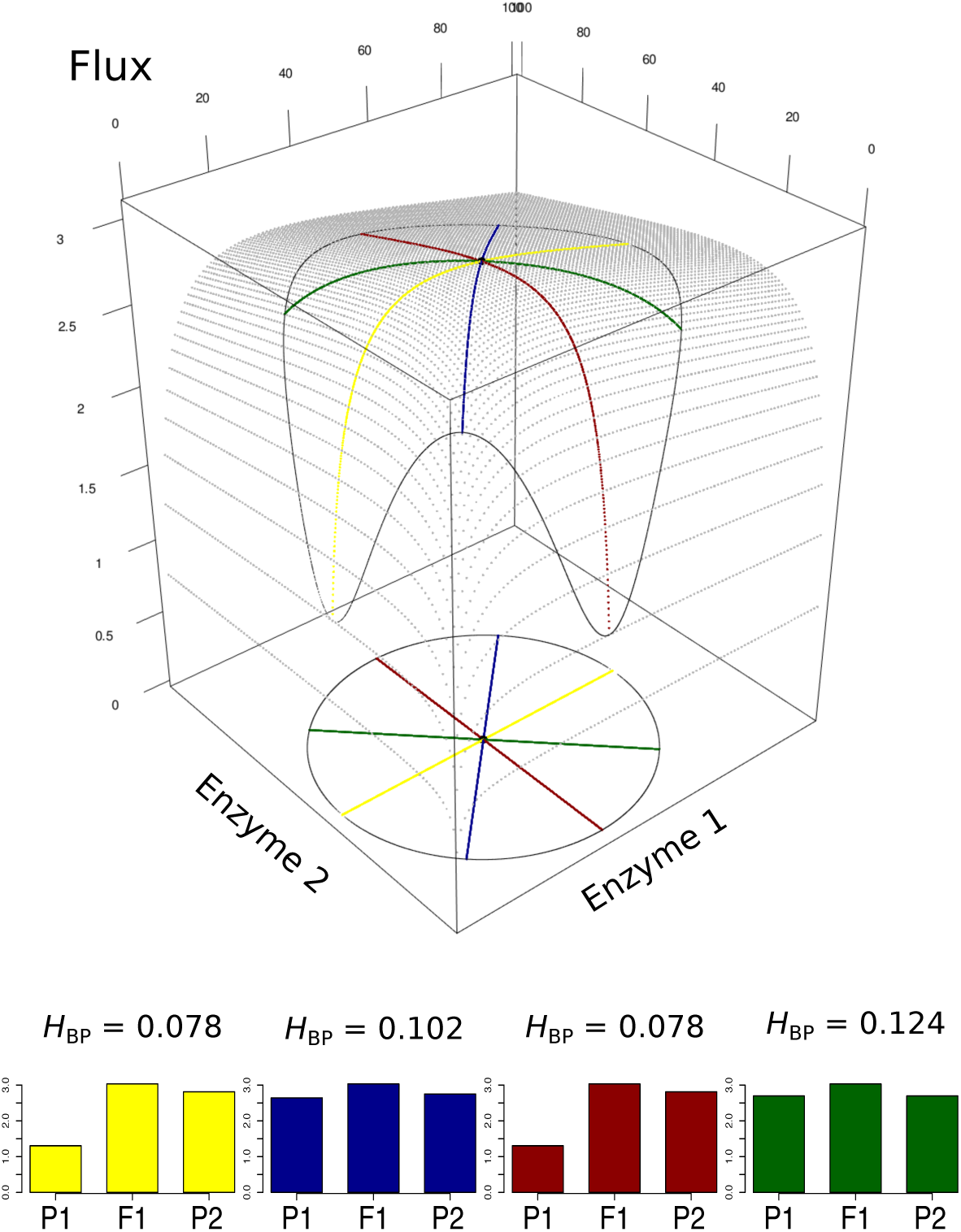
The geometry of heterosis for a constant enzymatic distance between parents, with constraint on *E*_tot_. Parameter values are the same as in Fig 14, but the constraint increases concavity, resulting in BPH in all cases. The flux equation is 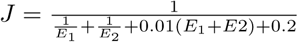

**S9 Fig.**
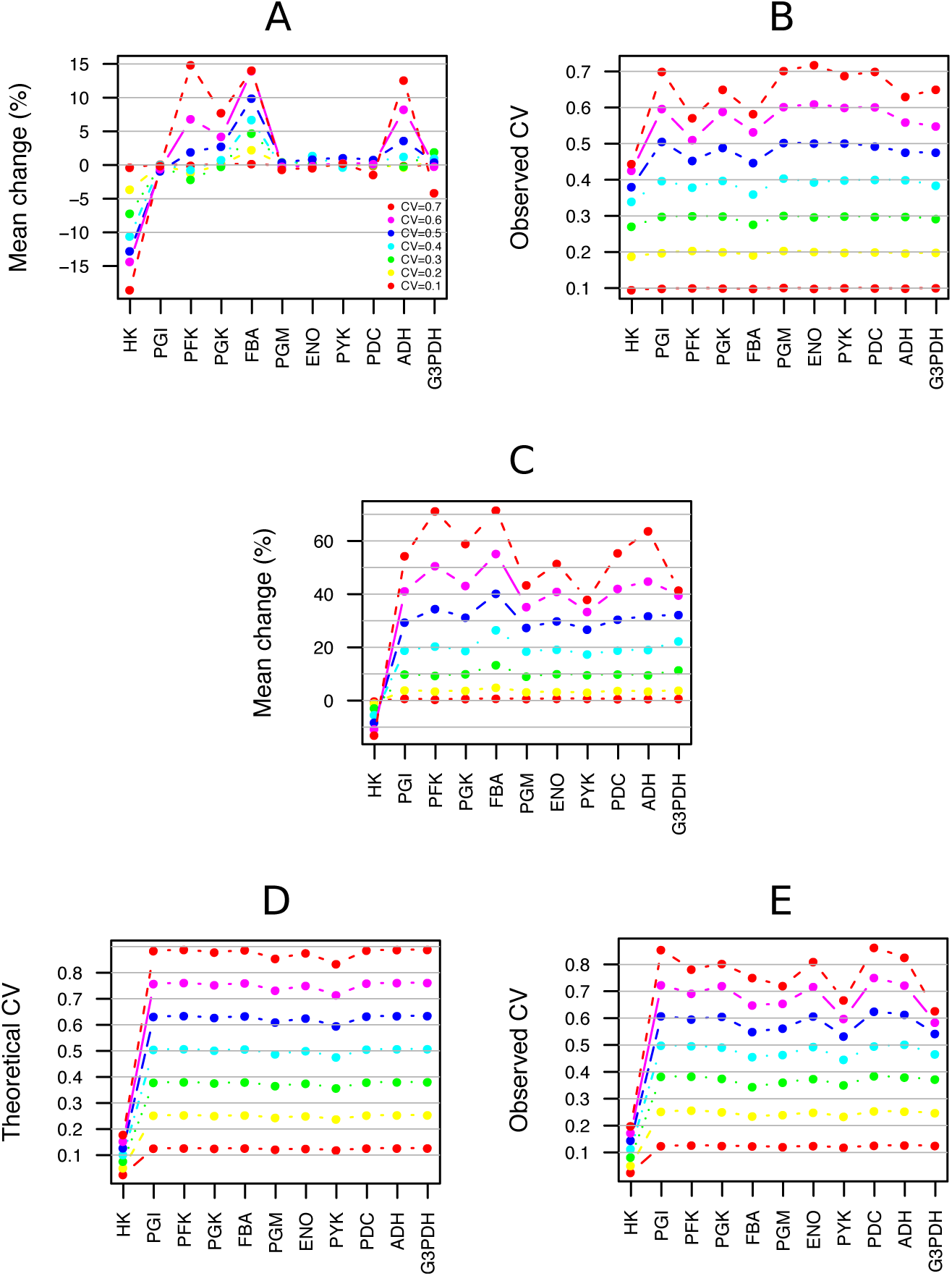
Deviations from the means and *c*_v_’s of enzyme concentrations relative to the initial values used in the simulations of the glycolysis/fermentation system. A and B: Free *E*_tot_. A: Variation in the means relative to the reference concentrations of the model (in %). B: Observed *c*_v_’s. C, D and E: Fixed *E*_tot_. C: Variation in the means relative to the reference concentrations of the model (in %). D: Theoretical *c*_v_’s when *E*_tot_ is constrained; *c*_v_’s are inversely proportional to the enzyme concentrations. E: Observed *c*_v_’s.

